# Hydroxysteroid 17-beta dehydrogenase 13 *(Hsd17b13)* knockdown attenuates liver steatosis in high-fat diet obese mice

**DOI:** 10.1101/2024.02.27.582262

**Authors:** Shehroz Mahmood, Nicola Morrice, Dawn Thompson, Sara Milanizadeh, Sophie Wilson, Philip D. Whitfield, George D. Mcilroy, Justin J. Rochford, Nimesh Mody

## Abstract

Hydroxysteroid 17-beta dehydrogenase 13 (*HSD17B13*) loss-of-function gene variants are associated with decreased risk of ‘metabolic dysfunction-associated steatotic liver disease’ (MASLD). Our RNA-seq analysis of steatotic liver from obese mice -/+ Fenretinide treatment identified major beneficial effects of Fenretinide on hepatic gene expression including *Hsd17b13*. We sought to determine the relationship between Hsd17b13 expression and MASLD and to validate it as a therapeutic target by liver-specific knockdown. Hsd17b13 expression, which is unique to hepatocytes and associated with the lipid-droplet, was elevated in multiple models of MASLD and normalised with prevention of obesity and steatotic liver. Direct, liver-specific, shRNA-mediated knockdown of *Hsd17b13* (*shHsd17b13*) in high-fat diet (HFD)-obese mice, markedly improved hepatic steatosis with no effect on body weight, adiposity or glycaemia. *shHsd17b13* decreased elevated serum ALT, serum FGF21 levels and markers of liver fibrosis e.g. *Timp2*. *shHsd17b13* knockdown in HFD-obese mice and Hsd17b13 overexpression in cells reciprocally regulated expression of lipid metabolism genes e.g. *Cd36*. Global lipidomic analysis of liver tissue revealed a major decrease in diacylglycerols (e.g. DAG 34:3) with *shHsd17b13* and an increase in phosphatidylcholines containing polyunsaturated fatty acids (PUFA) e.g. PC 34:3 and PC 42:10. Expression of key genes involved in phospholipid and PUFA metabolism e.g. *Cept1*, were also reciprocally regulated suggesting a potential mechanism of Hsd17b13 biological function and role in MASLD. In conclusion, *Hsd17b13* knockdown in HFD-obese adult mice was able to alleviate MASLD via regulation of fatty acid and phospholipid metabolism, thereby confirming HSD17B13 as a genuine therapeutic target for MASLD and development of liver fibrosis.

**KEY POINTS:**

- *HSD17B13* loss-of-function gene variants are associated with decreased risk of metabolic dysfunction-associated (MA) steatotic liver disease and steatohepatitis (MASLD and MASH).
- RNA-seq analysis of steatotic liver identified beneficial effects of Fenretinide on hepatic gene expression including downregulation of *Hsd17b13*.
- Liver-specific shRNA knockdown of *Hsd17b13* in obese mice markedly improved hepatic steatosis and markers of liver health e.g. serum ALT, serum Fgf21 levels.
- Hsd17b13 influenced expression of lipid/phospholipid metabolism genes e.g. Cd36 and Cept1 and phosphatidylcholines PC 34:3 and PC 42:10.
- Our study suggests a mechanism of HSD17B13’s biological function and the strong rationale behind targeting HSD17B13 for MASLD/MASH.

## INTRODUCTION

Metabolic dysfunction-associated steatotic liver disease (MASLD), previously termed non-alcoholic fatty liver disease (NAFLD) is a common complication associated with obesity and diabetes. MASLD can progress to the more severe metabolic dysfunction-associated steatohepatitis (MASH), previously termed non-alcoholic steatohepatitis (NASH) and therefore requires urgent therapeutic attention (Pafili & Roden, 2021). MASLD is characterised by an accumulation of triglycerides in the liver and is estimated to affect 25-30% of the global population as well as up to 90% of patients classed as morbidly obese (Pafili & Roden, 2021). Currently, there are no approved therapeutics available for MASLD other than dietary and lifestyle interventions, however human trials suggest medicines with known benefits against obesity and diabetes may be useful to also treat MASLD (Luo *et al.*, 2022). In addition, several novel pharmacologic therapies are in various phases of clinical development for the specific treatment of MASLD/MASH (Esler & Bence, 2019).

Several studies including human genome-wide association studies (GWAS) and multi-omics data analysis have identified genes that are key drivers of MASLD (Chella Krishnan *et al.*, 2018). For example, 17-beta hydroxysteroid dehydrogenase 13 (HSD17B13) is a novel lipid-droplet protein that has been identified from human steatotic liver biopsies (Su, W. *et al.*, 2014). Moreover, single nucleotide polymorphisms identified by GWAS linked to MASLD/MASH, performed in cohorts around the world, have identified several gene variants of *HSD17B13* (Amangurbanova *et al.*, 2023). Three independent gene variants in *HSD17B13* which generate loss-of-function proteins are reported to associate with protection from liver injury (Chella Krishnan *et al.*, 2018, Ma, Y. *et al.*, 2019, Abul-Husn *et al.*, 2018). HSD17B13 gene expression and protein levels are upregulated in patients and mouse models of MASLD and overexpression of Hsd17b13 in mice promotes rapid lipid accumulation in the liver of mice (Su, W. *et al.*, 2014). However, the traditional gene knockout of *Hsd17b13* in mice had no beneficial effect on the development of liver steatosis with various dietary interventions (Ma, Yanling *et al.*, 2021). This surprising result suggests that the normal biological role of HSD17B13 and its metabolite pathway maybe important for normal liver function from birth, but abnormally elevated levels of HSD17B13 in adulthood is a pathological driver of steatotic liver.

We have previously reported the beneficial effects of the synthetic retinoid Fenretinide to inhibit adiposity, insulin resistance and the accumulation of liver triglycerides (Mcilroy, G. D. *et al.*, 2013, Morrice *et al.*, 2017). In the present study, our analysis of unbiased, whole-genome RNA-sequencing in liver tissue from mice with high-fat diet (HFD) induced obesity and type-2 diabetes and mice treated with Fenretinide to inhibit MASLD (Mcilroy, G. D. *et al.*, 2013, Morrice *et al.*, 2017) revealed genes that are key drivers of MASLD including PPARα targets, lipid-droplet proteins e.g. *Hsd17b13*, pro-fibrotic genes, and phospholipid metabolism genes.

Hsd17b13 has been reported to have dehydrogenase activity towards retinol and thus modulate retinoic acid levels (Ma, Y. *et al.*, 2019). Thus, together with the human gene variant data on *HSD17B13* and liver disease, we were attracted to investigating this protein further. We hypothesised that direct inhibition of elevated levels of liver *Hsd17b13* using RNAi-mediated knockdown in vivo may reveal a beneficial phenotype and lead to novel insights into the biological function of this uncharacterised lipid-droplet protein.

## MATERIALS AND METHODS

### Animal studies

All animal procedures were performed under a project license approved by the U.K. Home Office under the Animals (Scientific Procedures) Act 1986 (PPL P1ECEB2B6) and the University of Aberdeen Ethics Review Board. Studies were performed following the recommendations in the ARRIVE guidelines under guidance by the Veterinary Surgeon and Animal Care and Welfare Officers of the institutional animal research facility. The Fenretinide HFD and *db/db* studies were previously described (summarized in Supplemental table S1) (Mcilroy, G. D. *et al.*, 2013, Morrice *et al.*, 2017). For the Hsd17b13 knockdown study, 3-4 week male C57BL/6J mice were purchased from Charles River. Mice were maintained at 22– 24°C on 12-h light/dark cycle with free access to food/water. At 11 weeks age, 18 mice were placed on to a 45% kcal HFD (Research diets, D12451) whilst 8 remained on regular Chow. Following 21 weeks on HFD, mice were randomised into two groups for intraperitoneal (i.p.) injection of AAV8-*shScrmbl* or AAV8-*shHsd17b13* (Vectorbuilder) at a virus titer of 1x10^11^ virus particles. Vector and shRNA information can be found at https://en.vectorbuilder.com id: VB180117-1020znr and id: VB200302-1251whe respectively. Chow controls received a saline injection of the same volume administered by i.p. injection. Body weight was measured weekly and physiological tests performed. Two weeks (14 days) after virus-shRNA injections, mice were humanely killed in accordance with guidelines and regulations detailed above, (following 5 hours fasting) by CO2-induced anaesthesia and cervical dislocation. Trunk blood following decapitation was collected and serum stored at -80°C. All tissues were rapidly dissected, frozen in liquid nitrogen, and stored at -80°C. Tissues for histology were fixed in formalin for 48h at 4°C, then stored at 4°C in phosphate-buffered saline (PBS) until further processing.

### Physiological tests

Body composition was measured before virus-shRNA injection to confirm HFD-induced obesity and aid with randomization into treatment/control groups. The measurement was repeated 7 days after shRNA treatment. Each mouse was analysed using an Echo MRI-3-in-1 scanner (Echo Medical Systems, Houston, TX, USA). Glucose tolerance test (GTT) was performed at 10 days post shRNA treatment. Briefly, mice were fasted for 5hrs, baseline glucose levels were sampled from tail blood using glucose meters (AlphaTRAK, Abbott Laboratories, Abbot Park, IL, USA). Subsequently mice were injected i.p. with 20% glucose (w/v) and blood glucose measured at 15, 30, 60 and 90 mins post-injection.

### Stable expression of HSD17B13 in cells

HEK293 cells and HepG2, were maintained in Dulbecco’s Modified Eagle Medium (DMEM) GlutaMAX with 10% foetal bovine serum (FBS) at 37 °C and 5% CO_2_. Human *HSD17B13* (NCBI Accession number: NM_178135.5) with HA-tag on the C-terminus was cloned into pcDNA3.1 (Genscript). HEK293 cells and HepG2 cells were transfected with HSD17B13 or empty control vector using Lipofectamine 2000, cultured for 2-3 weeks in G418 for neomycin resistance and positive colonies selected. Stable cell lines were treated with retinoic acid (RA, 1 μm, 2 h) or DMSO (vehicle control). Unsaturated/saturated fatty acid mixture (2:1 ratio of 0.66 mM oleic acid and palmitic acid 0.33 mM, 24 h) or 1% methanol (vehicle control) was prepared with 10% bovine serum albumin (fatty acid free) and 1% FBS in DMEM+G418.

### Immunoblotting

Frozen liver tissues were homogenised in 400µl of ice-cold Radioimmunoprecipitation assay buffer (10mM Tris-HCl pH 7.4, 150mM NaCl, 5mM EDTA pH 8.0, 1mM NaF, 0.1% SDS, 1% Triton X-100, 1% Sodium Deoxycholate with freshly added 1mM NaVO_4_ and protease inhibitors) using a PowerGen 125 homogeniser and lysates normalised to 1µg/µl. Proteins were separated on a 4-12% Bis-Tris gel (Fisher Scientific) by gel electrophoresis and transferred onto a nitrocellulose membrane. Membranes were probed with the following primary antibodies: HSD17B13 (1/500, ab122036, Abcam), GFP (1/1000, A11122, Invitrogen), GAPDH (1/1000, 14C10) and Vinculin (1/1000, 13901) both Cell Signaling Technology. Immunoblotting for RBP (Dako) was from 1µl serum (Yang, Q. *et al.*, 2005).

### RNA extraction and RT-qPCR

Frozen tissues were lysed in TRIzol reagent (Sigma) and RNA isolated using phenol/chloroform extraction according to manufacturer’s instructions. RNA was then synthesized into cDNA Tetrokit (Bioline, UK) and subjected to qPCR analysis using SYBR green and LightCycler 480 (Roche). Gene expression was determined relative to the reference gene *Nono* for mouse liver tissue and *Hprt* for human cell lines (HEK and HepG2) and analysis was performed using the Pfaffl method (Pfaffl, 2001). Primer sequences can be found in Supplemental table S2.

### Liver Histology

Liver tissues were embedded in paraffin, sectioned and stained with haematoxylin and eosin (H&E). Images were taken using a light microscope, EVOS XL (Thermofisher Scientfic, UK) at 20x and 40x magnification. Frozen liver tissue was used for fluorescent staining. Briefly, processed slides were washed PBS for 10-12 min. Subsequently, slides were incubated with Blocking (5% goat serum in PBS) and then with HCS Lipidtox^TM^ deep red neutral lipid stain (H34477) (1:200 in PBS) in RT each for 1 h. Furthermore, they were mounted with fluoroshield mounting medium with DAPI (ab104139).

### Liver and serum assays

50-100mg of frozen liver tissues were homogenised in 1ml of PBS and frozen in liquid nitrogen to enable further cell lysis. Samples were left to thaw at room temperature and centrifuged for 15s at 7500rpm to pellet debris. The remaining fat cake was resuspended in PBS, the resulting supernatant used to determine total liver triglycerides according to manufacturer’s instructions (Sigma, cat: MAK266). Serum was analysed for alanine aminotransferase (ALT) activity (Abcam, ab105134), FGF21 (R&D Systems, #MF2100) and total triglycerides (Sigma, cat: MAK266) following manufacturer’s guidelines.

### Lipidomics Analysis

Global lipidomics analysis was performed by high resolution liquid chromatography-mass spectrometry (LC-MS) using an Exactive Orbitrap mass spectrometer (Thermo Scientific, Hemel Hempsted, UK) interfaced to a Thermo UltiMate 3000 RSLC system. Samples (10 μl) were injected onto a Thermo Hypersil Gold C18 column (1.9 μm; 2.1 mm x 100 mm) maintained at 50°C. Mobile phase A consisted of water containing 10 mM ammonium formate and 0.1% (v/v) formic acid. Mobile phase B consisted of a 90:10 mixture of isopropanol-acetonitrile containing 10 mM ammonium formate and 0.1% (v/v) formic acid. The initial conditions for analysis were 65%A-35%B, increasing to 65%B over 4 min and then to 100%B over 15 min, held for 2 min prior to re-equilibration to the starting conditions over 6 min. The flow rate was 400 μl/min. Analyses were undertaken in positive and negative ion modes at a resolution of 100,000 over the mass-to-charge ratio (m/z) range of 250 to 2,000. Progenesis QI v3.0 (Nonlinear Dynamics, Newcastle upon Tyne, UK) was used to process the data sets and lipids were identified through interrogation of HMDB (http://www.hmdb.ca/), LIPID MAPS (www.lipidmaps.org/). Multivariate statistical analysis was performed using SIMCA-P v13.0 (Umetrics, Umea, Sweden) and the heatmap was generated with the online available platform MetaboAnalyst 5.0 (https://www.metaboanalyst.ca/) (Pang *et al.*, 2021).

### RNA sequencing

As previously described (Morrice *et al.*, 2017). Total RNA was extracted from frozen liver using TriZOL reagent (Sigma Aldrich, Irvine, UK) to manufacturer’s instructions. RNA was purified using RNeasy miniprep kit (Qiagen, Manchester, UK) and quantified on a Bioanalyzer 2000 (Agilent Technologies, Edinburgh, UK). Ribosomal RNA was removed from samples using the Ribo-Zero rRNA removal kit (Illumina, San Diego, CA, USA). Sequencing libraries were prepared using the TruSeq RNA Library Preparation Kit v2 (Illumina) following the low sample protocol, using 1 μg RNA. Libraries were sequenced on the NextSeq-500 Desktop sequencer platform (Illumina) with a 2 × 75 paired-end read length giving 719 million raw reads (average depth 45 million reads per sample). Reads were filtered using the FASTQ Toolkit v2.0.0 in the Illumina BaseSpace cloud computing environment to remove adapter sequences and reads with phred score <20 or length <20 bp. Filtered reads were checked using FastQC before aligning with STAR 2.0 to the *Mus musculus* UCSC mm10 genome and differential expression measured using DESeq2 with default parameters. RNA-seq data generated in this study has been deposited in the NCBI Gene Expression Omnibus (GEO) database accession number GSE220684.

### Statistical Analysis

All values are expressed as mean + S.E.M, unless otherwise stated. Statistical analyses were performed by using one- or two-way ANOVA (where stated) followed by multiple-comparison tests to compare the means of three or more groups. In all figures, */^#^p<0.05, **/^##^p<0.01, ***/^###^p≤0.001. All analyses were performed using GraphPad Prism (GraphPad Software).

## RESULTS

### Hepatic RNA-seq reveals signature gene expression alterations in HFD-induced obese steatotic liver and rescue by beneficial effects of Fenretinide

We have previously reported the effects of Fenretinide to inhibit adiposity, insulin resistance and the accumulation of liver triglycerides (summarized in Supplemental Table S1). In a **novel** approach to characterise the molecular mechanism of beneficial effects of Fenretinide in the liver, we performed RNA-seq (GEO number GSE220684) on liver tissue from HFD-induced obese C57BL/6J mice +/- Fenretinide for 20 weeks, compared to lean control mice (Figure 1A, Supplemental Figure S1) (Mcilroy, G. D. *et al.*, 2013, Morrice *et al.*, 2017). HFD-induced obesity resulted in upregulation of 941 liver genes and downregulation of 637 genes (Figure 1B). Many of the genes upregulated in HFD steatotic liver were involved in triglyceride synthesis and fatty acid metabolism and well-established targets of PPARα signalling and/or GWAS identified genes associated with MASLD (e.g *Mogat, Agpat9, Crat and Vnn1, Hsd17b13, Pnpla3* and *Inhbe* Figure 1C, Supplemental Figure S1) (Kersten, 2014). Fenretinide treatment partially prevented (and in some cases completely normalised) the increase in expression of many of these genes (Figure 1C).

**Figure 1:**
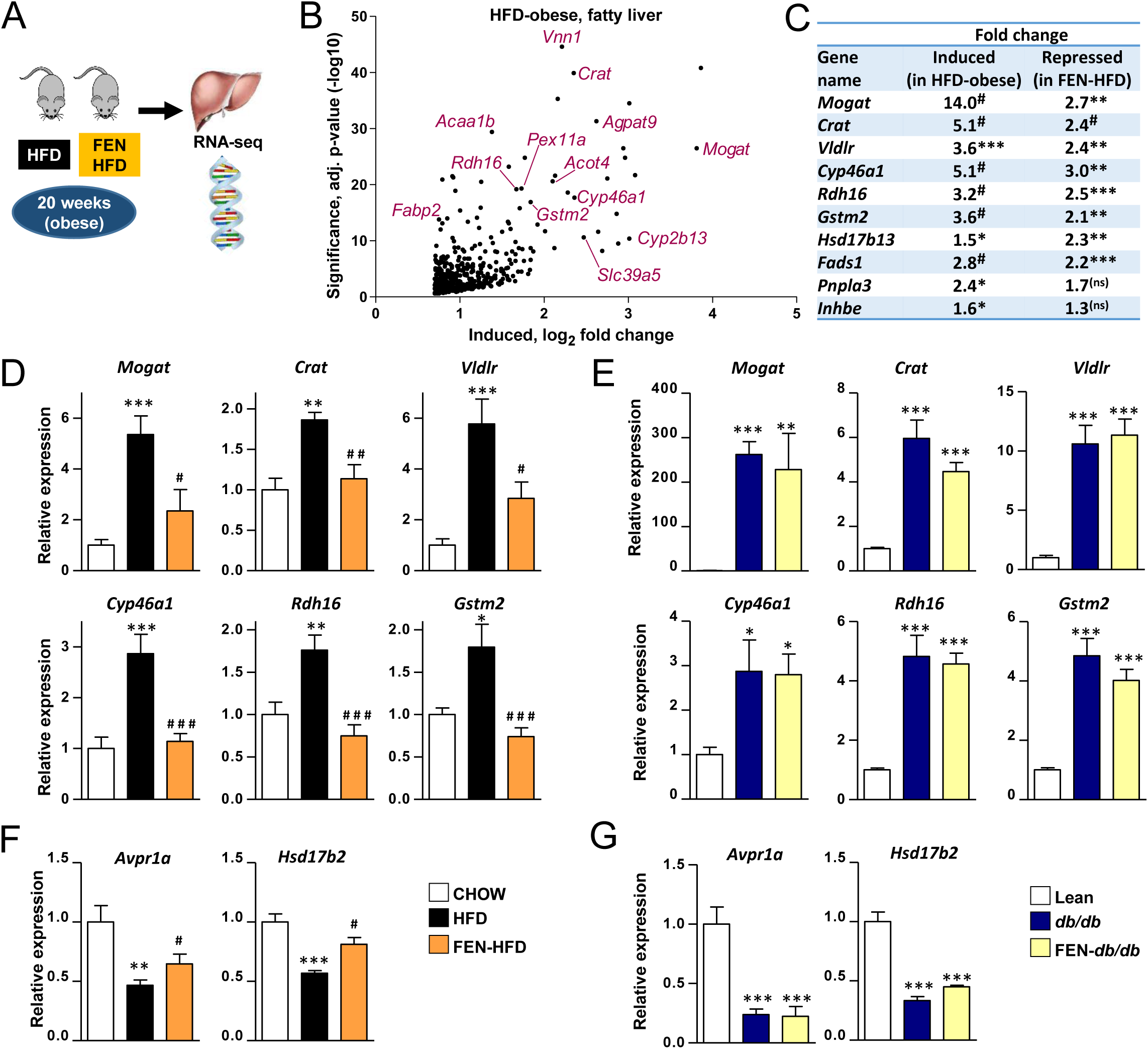
Hepatic gene expression in high-fat diet-induced obese and genetically-obese mice with liver steatosis, and effects of Fenretinide treatment. **A.** Graphical illustration of mouse liver study described. **B.** Volcano plot of hepatic RNA-seq data of HFD (obese-steatotic liver) most highly induced genes by fold change and significance, compared to lean control mice (Morrice *et al.*, 2017). **C.** Shortlist of differential gene expression from RNA-seq data, reciprocally regulated genes in HFD (obese-steatotic liver) +/-FEN, involved in triglyceride synthesis and fatty acid metabolism and genes associated with MASLD. HFD induced genes vs lean control mice and FEN-HFD repressed genes vs HFD, significance, adjusted p-value *** <10^-15^ , ** <10^-8^ , * <10^-3^ , ns not significant. **D.** and **F**. Hepatic gene expression in HFD (obese-steatotic liver) mice +/-FEN. **E.** and **G**. Hepatic gene expression in *db/db* (genetically-obese steatotic liver) mice +/- FEN. Data (n=7-8 per group) are presented as mean + SEM and analysed by one-way ANOVA followed by Bonferroni multiple comparison tests where *p≤0.05, **p≤0.01 and ***p≤0.001 (HFD vs CHOW and *db/db* vs lean) or # p≤0.05 and ## p≤0.01 and ### p≤0.001 (FEN-HFD vs HFD and FEN-*db/db* vs *db/db*).

HFD in the C57BL/6J strain is typically reported not to induce fibrosis or inflammation and therefore other diets are utilized to induce hepatic injury (for example GAN diet or choline-deficient HFD) (Loft *et al.*, 2021, Vacca *et al.*, 2024). However, our RNA-seq revealed that HFD-induced obese in C57BL/6J mice for 20 weeks did induce increased expression of genes driving fibrosis and tissues remodelling e.g. collagens (*Col1a1*, *Col12a1* and *Col15a1*), tissue inhibitors of matrix metalloproteinases (*Timp1* and *Timp2*) and canonical Yap/Taz signalling pathway (*Wwtr1* and *Ctgf*) (Supplemental Figure S1). Moreover, Fenretinide treatment prevented the increase in expression of many of these genes (Supplemental Figure S1).

We were able to confirm the differential gene expression changes by RNA-seq in both chronic HFD-induced obese groups (+/- Fenretinide, 20 weeks) against two lean control groups (HFD +/- Fenretinide, 7 days) where we determined similar results with induction of PPARα targets, lipid-droplet proteins, pro-fibrotic genes, and phospholipid metabolism genes (Supplemental Figure S1). However, the relative level of induction by 20 weeks HFD and reciprocal inhibition by Fenretinide differed widely between genes. The acute effects of Fenretinide treatment (7 days) is not the focus of this present study (manuscript in preparation).

To further characterise the effects on differential gene expression changes linked to steatotic liver, we compared the effects of Fenretinide on hepatic gene expression in HFD- induced and genetically obese *db/db* mouse model (Morrice *et al.*, 2017). Both HFD and *db/db* obese mice had increased expression of hepatic lipid metabolism genes (e.g. *Mogat* and *Crat*) (Figure 1D and 1E). Fenretinide treatment prevented this increase in HFD mice but had no effect in *db/db* mice (Figure 1D and 1E). Similarly, HFD repressed other genes (e.g. *Avpr1a* and *Hsd17b2*) and Fenretinide treatment prevented this decrease in HFD mice but had no effect in *db/db* mice (Figure 1F and 1G). These novel findings regarding the mechanism of Fenretinide action suggests that the beneficial effects on hepatic gene expression are associated with inhibition of adiposity (induced by HFD feeding) and associated insulin resistance and MASLD but not with Fenretinide treatment in genetically-obese *db/db* where adiposity and MASLD were not inhibited by this synthetic retinoid (Mcilroy, G. D. *et al.*, 2013, Morrice *et al.*, 2017).

### Hsd17b13 expression is upregulated in HFD-induced and genetic mouse models of obesity, hyperglycaemia and MASLD

17β-Hydroxysteroid dehydrogenases (17β-HSDs) are a family of 15 members, which play a key role in sex hormone metabolism by catalyzing steps of steroid biosynthesis (e.g. Hsd17b1 and b2) (Poutanen & Penning, 2019, Marchais-Oberwinkler *et al.*, 2011). Some may also play a role in cholesterol and fatty acid metabolism. Of the hydroxysteroid 17-beta dehydrogenase family of genes the most closely related to *Hsd17b13* is *Hsd17b11* although it is not specific to hepatocytes and also expressed in adipocytes (Horiguchi *et al.*, 2008). HFD- induced obesity also upregulated *Hsd17b11 and Hsd17b7* hepatic gene expression in C57BL/6 mice and Fenretinide treatment decreased their gene expression compared to HFD mice (new Figure 2A RNA-seq mini-table Hsd17b family of genes). HFD also upregulated *Hsd17b4* and *Hsd17b12* hepatic gene expression in C57BL/6 mice, however, Fenretinide treatment had no effect to decrease these genes. *Hsd17b13* has been linked to MASLD and was one of the top 10% most upregulated genes in C57BL/6 HFD mice, increased 1.5-fold (Figure 2A and 2B).

**Figure 2:**
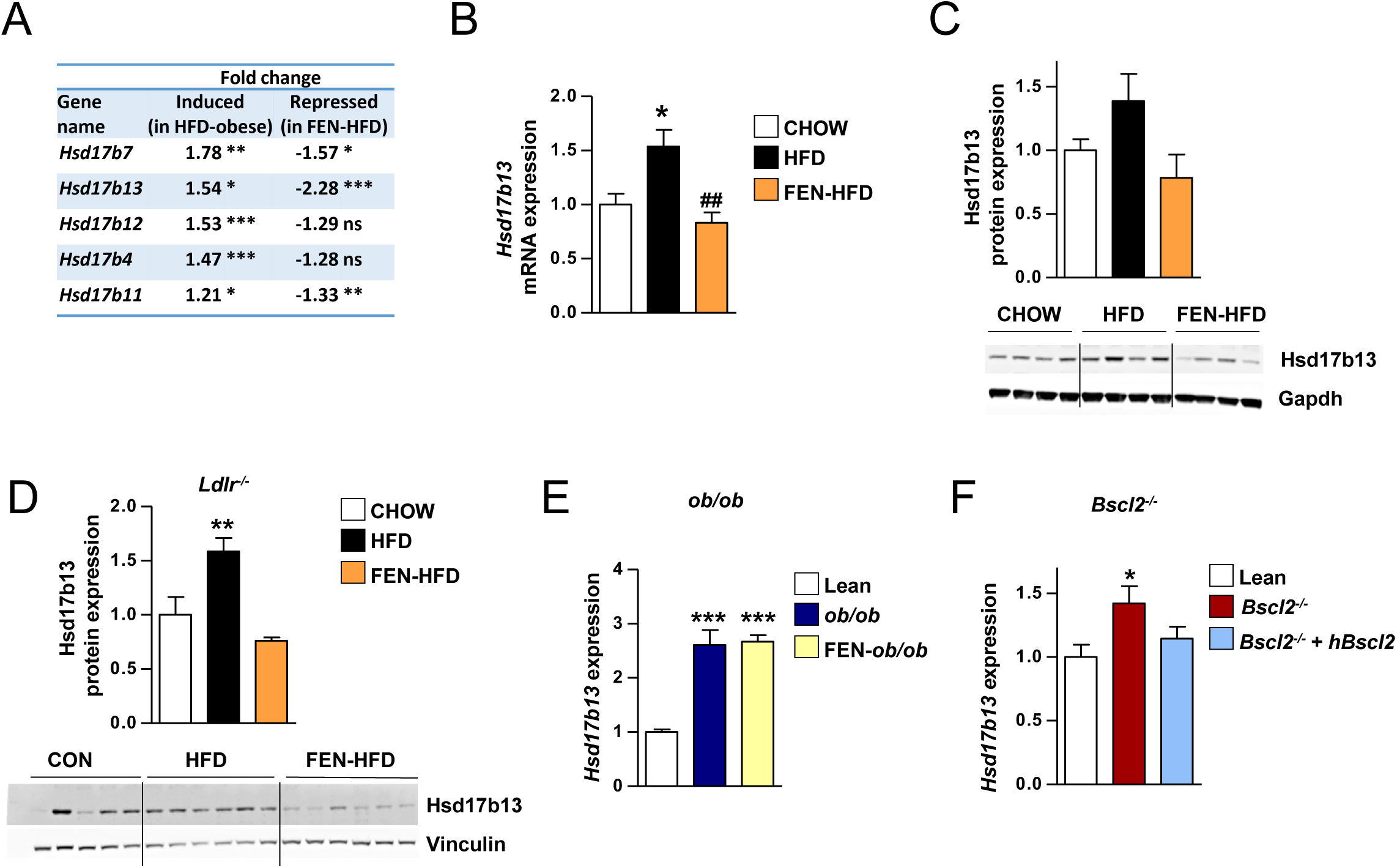
Increased Hsd17b13 expression in a HFD-induced and genetic mouse models of obesity, hyperglycaemia and MASLD. **A.** Differential gene expression from RNA-seq data. Hydroxysteroid 17-beta dehydrogenase family of genes reciprocally regulated in HFD (obese-steatotic liver) +/-FEN. HFD induced genes vs lean control mice and FEN-HFD repressed genes vs HFD, significance, adjusted p-value # <10^-15^ , *** <10^-8^ , ** <10^-3^ , * <0.05 ns not significant. **B.** *Hsd17b13* gene expression in HFD (obese-steatotic liver) mice +/-FEN and chow fed lean controls, normalised to *Nono*, (n=7-8 per group). **C.** Western blot of Hsd17b13 protein expression in HFD (obese-steatotic liver) mice +/-FEN and chow fed lean controls, and quantification normalised to Gapdh, (n=4 per group). **D.** Western blot and quantification of HSD17B13 protein expression in *Ldlr^-/-^* HFD (obese- steatotic liver) mice +/-FEN and control (10% kcal) fat diet lean controls, and quantification normalised to vinculin, (n=5-6 per group). **E.** *Hsd17b13* gene expression in *ob/ob* (genetically-obese steatotic liver) mice +/-FEN and control (10% kcal) fat diet lean controls, normalised to *Nono*, (n=6-7 per group). **F.** *Hsd17b13* gene expression in *Bscl2*^-/-^ (lipodystrophic steatotic liver) mice +/- human *Bscl2*^-/-^ rescue and wildtype control, normalised to *Nono*, (n=8-9 per *Bscl2*^-/-^ groups, n=11 wildtype control). Data are presented as mean + SEM and analysed by one-way ANOVA followed by Bonferroni multiple comparison tests where *p≤0.05, **p≤0.01 and ***p≤0.001 (respective MASLD model vs control) or ## p≤0.01 (FEN-HFD vs HFD).

Since Hsd17b13 has been reported to have dehydrogenase activity towards retinol (Ma, Y. *et al.*, 2019), was one of the most repressed genes with retinoid treatment and together with the human gene variant data on Hsd17b13, this warranted further examination of this hepatocyte-specific 17β-HSD family member.

Hsd17b13 hepatic protein expression was also induced by a similar amount in HFD mice compared to lean controls (Figure 2C). Fenretinide treatment was able to prevent this induction, with both gene and protein expression levels close to lean C57BL/6 control mice (Figure 2A, 2B and 2C). Fenretinide treatment also repressed Hsd17b13 expression in parallel with prevention of liver steatosis in LDLR^-/-^ mice fed-HFD plus high-cholesterol diet model of dyslipidemia (Figure 2D) (Thompson *et al.*, 2023). *Hsd17b13* gene expression levels were induced in two more genetical models of metabolic disease and steatotic liver, in leptin-deficient obese *ob/ob* mice (Figure 2E) and in lipodystrophic Seipin/*Bscl2*^-/-^ mice (Figure 2F) (Mcilroy, George D. *et al.*, 2018). FEN treatment could not inhibit the increase in *Hsd17b13* obese *ob/ob* mice but gene therapy to restore WAT in *Bscl2*^-/-^ mice inhibited MASLD and *Hsd17b13* gene expression (Figure 2F) (Sommer *et al.*, 2022). These data and taken together with the human gene variant data suggest a role of 17β-hydroxysteroid dehydrogenases in liver lipid metabolism and an important role for Hsd17b13 in MASLD (Chella Krishnan *et al.*, 2018, Ma, Y. *et al.*, 2019, Abul-Husn *et al.*, 2018).

### RNAi-mediated Hsd17b13 knockdown in HFD-induced obesity and MASLD does not affect body weight, adiposity or glucose homeostasis

This RNAi-mediated knockdown translational strategy has been taken by multiple pharma/biotech in on-going clinical trials (Amangurbanova *et al.*, 2023). Candidate short hairpin RNA (shRNA) were tested for knockdown efficiency in cells overexpressing mouse *Hsd17b13*, and the most potent (hereafter renamed *shHsd17b13*) was selected for treatment *in vivo* (Supplementary Figure S2). To evaluate the role of Hsd17b13 in driving steatotic liver, we proceeded to directly knockdown the elevated levels of this protein in adult mice with HFD- induced MASLD thereby circumventing the lack of a protective effect of Hsd17b13 whole-body, total genetic knockout from birth in mice. We did not utilize a specific diet (e.g. GAN diet or choline-deficient HFD) to induce fibrosis or inflammation since these tend to suppress weight gain and adiposity, the primary drivers of MASLD) (Loft *et al.*, 2021, Vacca *et al.*, 2024, Luukkonen *et al.*, 2023). Moreover, we had already determined the reciprocal regulation of *Hsd17b13* expression was associated with a simple HFD-induced obesity +/- beneficial effects of Fenretinide (Figure 1 and 2).

HFD caused increased weight gain and obesity over time compared to chow-fed C57BL/6 mice, as expected (Figure 3A). At 21 weeks on HFD, obese mice were randomised for administration of adeno-associated virus (AAV)-8 constructs encoding either *shHsd17b13* or scrambled control (*shScrmbl*) with the U6 promoter for high expression of shRNAs. *shHsd17b13* treatment had no effect on body weight or adiposity compared to HFD-*shScrmbl* control in the following 2 weeks (Figure 3B and C). Hepatic GFP co-expression confirmed successful AAV8 delivery in HFD mice (Figure 3D). *shHsd17b13* treatment suppressed the elevated *Hsd17b13* expression compared to *shScrmbl* treatment, normalising gene and protein expression to a level similar to that in chow-fed lean mice (Figure 3E and F). HFD-induced obesity caused glucose intolerance and *shHsd17b13* knockdown had no effect to alter this compared to *shScrmbl* control mice (Figure 3G, H and I).

**Figure 3:**
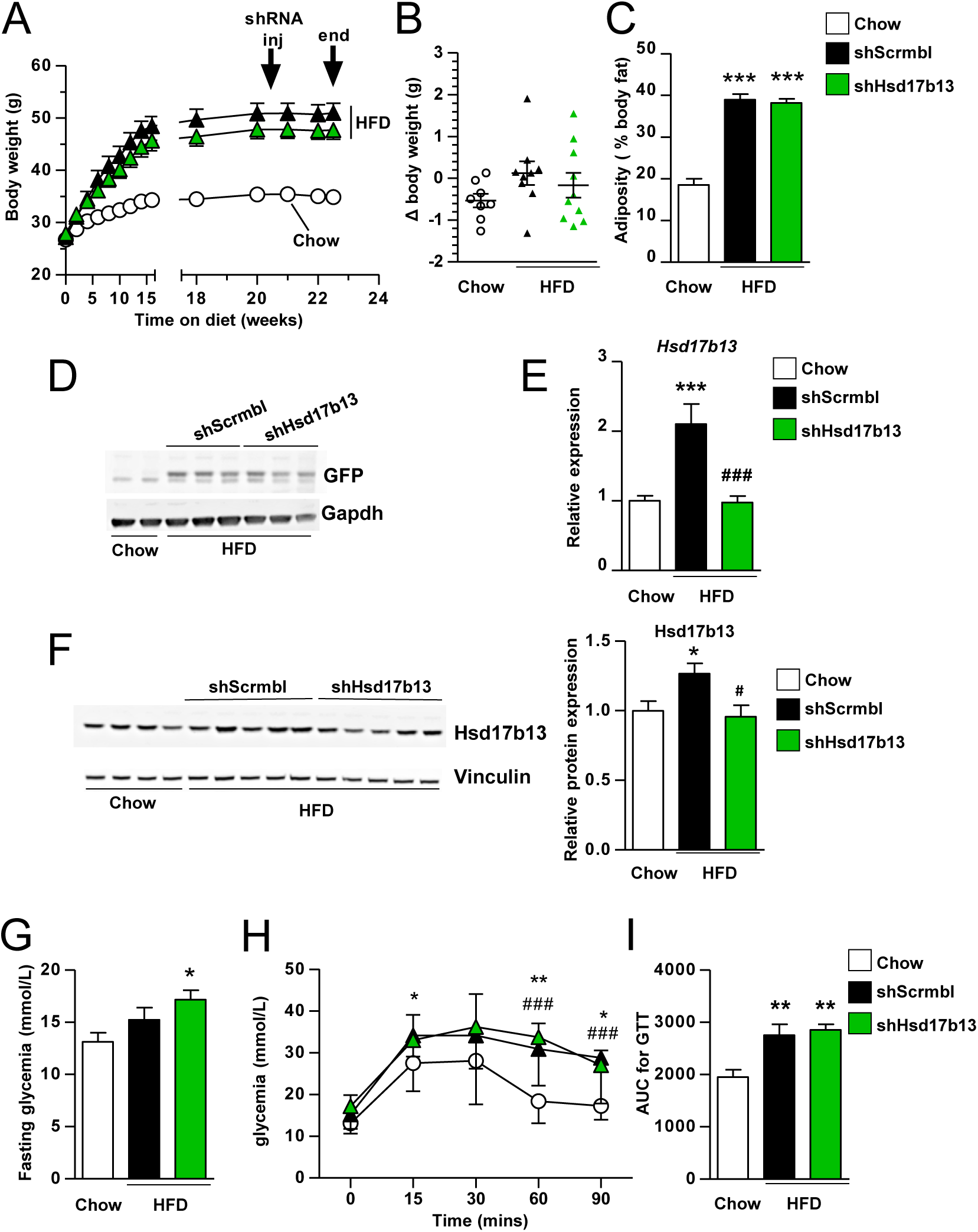
Effect of RNAi-mediated Hsd17b13 knockdown on HFD-induced obesity and glucose homeostasis. **A.** Body weights of C57BL/6 mice on high-fat diet (HFD, 45% kCal fat) or control CHOW diet for 23 weeks. At 21 weeks on HFD, obese mice were randomised for administration of AAV8- GFP-*shHsd17b13* or scrambled control shRNA (*shScrmbl*), n=8-9 per group. **B.** Change in body weight from 21 to 23 weeks diet and **C.** Adiposity (% body fat) at week 22 HFD in response to administration of AAV8-GFP-shRNA. At 23 weeks, following administration of AAV8-GFP-*shHsd17b13* or *shScrmbl*, **D.** Western blot of GFP in representative mouse liver tissues **E.** *Hsd17b13* gene expression in mouse livers, n=8 per group **F.** Western blot of Hsd17b13 protein expression and vinculin (loading control) in mouse livers and quantification, n=4-5 per group. **G.** Fasting (5 hours) glycaemia (basal glycemia from GTT). *p≤0.05 (*shHsd17b13* HFD vs Chow) **H.** Glucose tolerance test (GTT) performed in week 2 post-treatment with *shHsd17b13* or *shScrmbl*, n=8-9 per group, **I.** Area under curve (AUC) from GTT for each individual mouse. All data presented as mean ± SEM and analysed by one-way ANOVA followed by Bonferroni multiple comparison tests where *p≤0.05 , **p≤0.01 and ***p≤0.001 (HFD groups vs Chow) #p≤0.05 and ### p≤0.001 (HFD- *shHsd17b13* vs HFD-*shScrmbl*) In addition, in **F**. Western blot quantification, One-way ANOVA #p≤0.05 (HFD- *shHsd17b13* vs HFD-*shScrmbl*) but required t-test for *p≤0.05, (HFD-*shScrmbl* vs Chow). in **H**. GTT, t-tests were performed at individual time-points and where ### p≤0.001 (HFD-*shHsd17b13* vs Chow).

### RNAi-mediated Hsd17b13 knockdown prevents HFD-induced MASLD and markers of fibrotic injury

HFD-induced obesity and glucose intolerance led to steatotic liver characterised by increased number and size of lipid droplets as determined by H&E staining of liver tissues and fluorescent staining of lipid droplets (Figure 4A and B). Strikingly, *Hsd17b13* knockdown substantially decreased the number and size of hepatic lipid droplets in HFD mice. In addition, *shHsd17b13* decreased liver triglycerides by 45% from the elevated level in HFD-*shScrmbl* and back towards a normal level in CHOW mice (Figure 4C *first panel*). HFD-induced liver steatosis was associated with elevated levels of serum ALT and serum Fgf21 compared to lean mice (Figure 4C *second and third panels*). *shHsd17b13* decreased both serum ALT levels and serum Fgf21 indicating biomarkers of improved liver health compared to HFD-*shScrmbl* mice.

**Figure 4:**
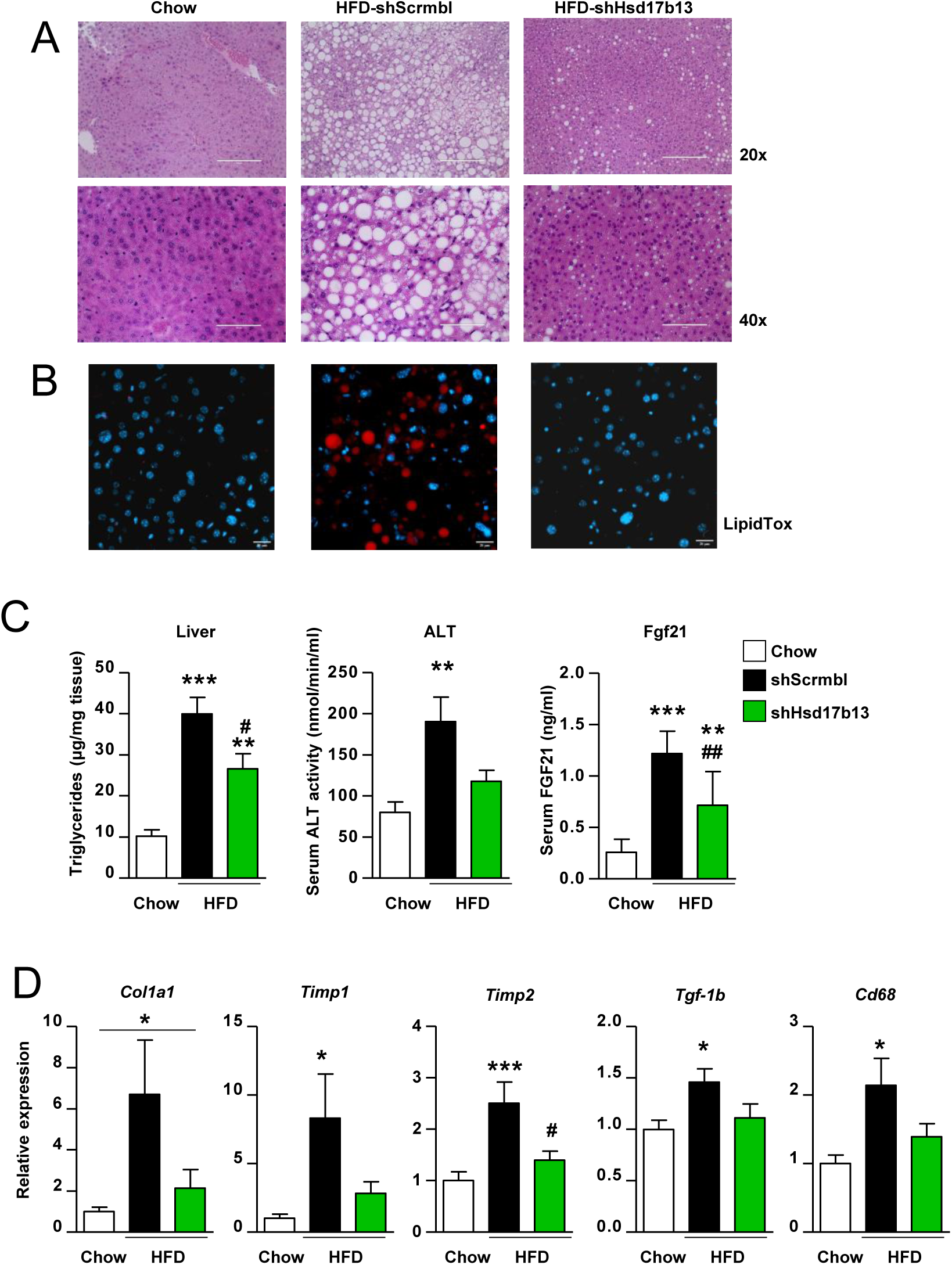
Effect of RNAi-mediated Hsd17b13 knockdown on HFD-induced MASLD. **A.** H&E staining of representative liver tissues at 20x and 40x magnification and **B.** Fluorescent staining of lipid droplets in C57BL/6 control CHOW diet mice or HFD obese mice treated with *shHsd17b13* or scrambled control (*shScrmbl*). Scale bars are 200 μm (20x) and 100 μm (40x) in H&E images and 20 μm in fluorescent images respectively. **C.** Liver triglyceride levels, serum ALT activity and serum FGF21 levels. **D.** Gene expression of fibrosis markers in liver tissue, Collagen type I alpha 1 (*Col1a1*), Collagen type IV alpha 1 (*Col4a1*), Tissue inhibitor of metalloproteinases (*Timp*)-*1* and *Timp2*, Transforming growth factor beta (*Tgf-*β)-1 and Cluster of Differentiation (*Cd*)-*68*. Data in **C**) and **D**) are presented as mean ± SEM (n=8 per group) and analysed by one-way ANOVA followed by Bonferroni multiple comparison tests where * p≤0.05, ** p≤0.01 and *** p≤0.001 (comparing HFD groups to control CHOW diet) or # p≤0.05 and ## p≤0.01 (comparing HFD-*shHsd17b13* to HFD-*shScrmbl*).

Persistent excess lipid accumulation and the progression to MASH, is associated with activation of hepatic stellate cells by TGF-β which leads to the development of fibrosis by collagens and matrix remodeling by tissue inhibitors of matrix metalloproteinases (Friedman *et al.*, 2018). Similar to the RNA-seq HFD +/- Fenretinide study (Figure 1 and Supplemental Figure S1), a 23 weeks HFD increased the expression of *Col1a1*, *Timp1*, *Timp2*, *Tgf-*β (Figure 4D).

Hsd17b13 knockdown significantly decreased expression of *Timp2* and trended to decrease expression of *Col1a1*, *Col4a1* and *Timp1, Tgf-*β in HFD mice. Similar changes in the pro-inflammatory macrophage marker *Cd68* were determined (individual data points for mice are in Supplemental Figure S3). Thus, *Hsd17b13* knockdown and attenuation of excess hepatic triglycerides storage, appeared to inhibit the development of fibrosis and MASH, via suppressing gene expression alterations associated with hepatic stellate cell activation and a pro-inflammatory environment.

### Increasing or decreasing Hsd17b13 levels does not affect retinoid transcriptional regulator genes or serum RBP4

Hsd17b13 has been reported to have *in vitro* dehydrogenase activity towards retinol and thus modulate retinoic acid levels in cells transfected with *Hsd17b13* (Ma, Y. *et al.*, 2019). Thus, we determined whether targets of retinoic acid signalling were altered with Hsd17b13 knockdown. HFD increased hepatic *Lrat* expression which is associated with altered retinyl ester dynamics in stellate cells and MASH (Figure 5A) (Molenaar *et al.*, 2020, Saeed *et al.*, 2017).

**Figure 5:**
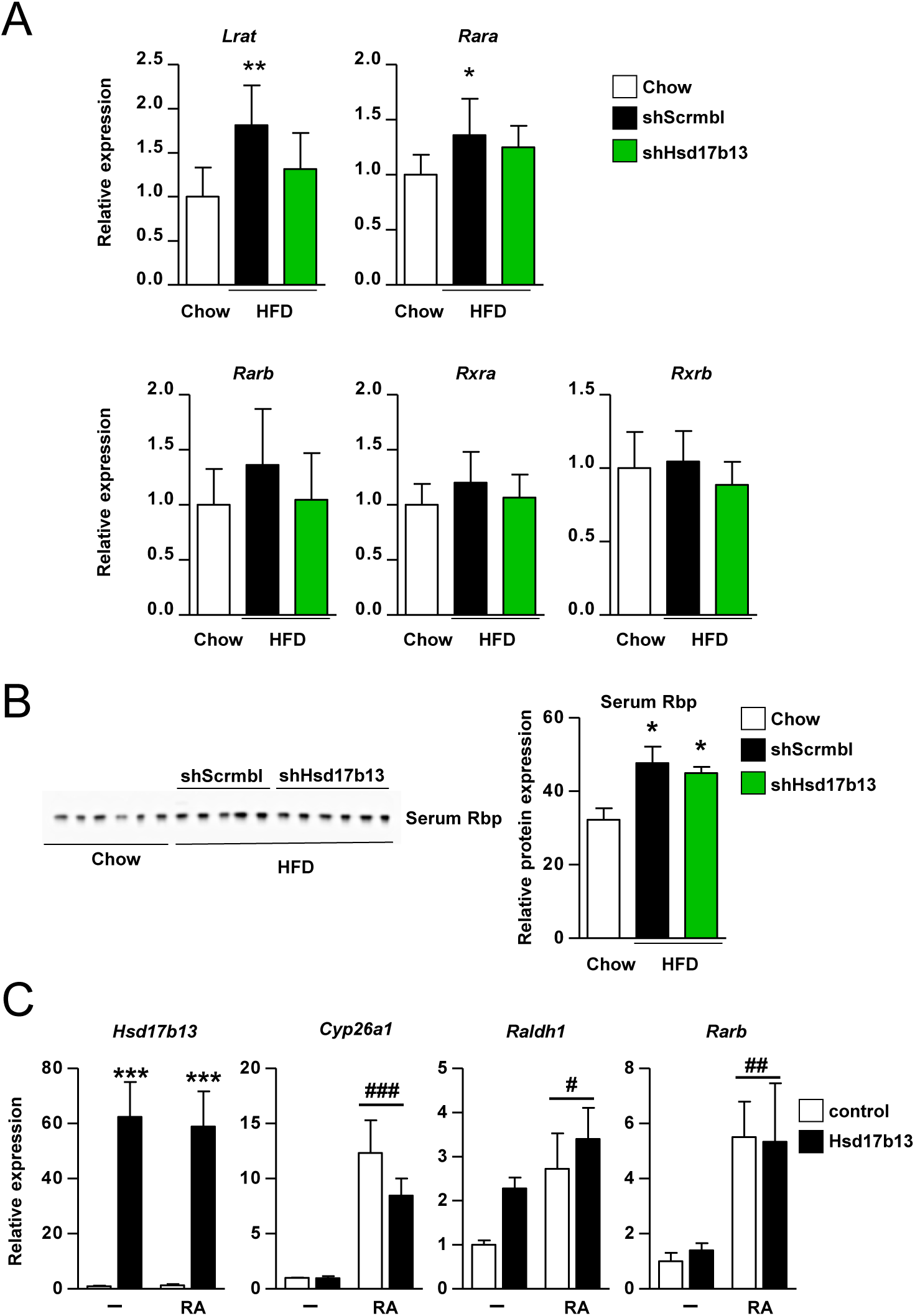
Effect of altered Hsd17b13 levels on retinoid genes and serum RBP4. **A.** Gene expression of hepatic lecithin-retinol acyltransferase (*Lrat*) and *Retinoic Acid* and *Retinoid X receptors* (*Rar* and *Rxr*) transcription factors (n=8 per group) in C57BL/6 control CHOW diet mice or HFD obese mice treated with *shHsd17b13* or scrambled control (*shScrmbl*). **B.** Western blot and quantification of serum RBP4 levels (n=5-6 per group). Data is presented as mean ± SEM with analysed by One-way ANOVA (*p≤0.05) followed by Bonferroni multiple comparison tests where * p≤0.05 and ** p≤0.01 (comparing HFD groups to control CHOW diet). **C.** HEK293 cells were transfected with HA-tagged Hsd17b13 or pcDNA3.1 control plasmid to stably express Hsd17b13 and treated with retinoic acid (RA, 1 μm, 2 h) or DMSO (vehicle control). Gene expression of *Hsd17b13* and retinoic acid metabolism genes *Cyp26a1*, *Raldh1* and *Rarb*. Data is presented as mean ± SEM from n=3 biological replicates and analysed by two-way ANOVA where *** p≤0.001 (Hsd17b13 cells vs pcDNA control cells), # p≤0.05 and ## p≤0.01 and ### p≤0.001 (RA vs DMSO control). Hsd17b13 over-expression did increase *Raldh1* expression by >2-fold (p<0.001 by t-test)

Hsd17b13 knockdown trended to decrease expression of *Lrat. shHsd17b13* had no effect to alter the expression of nuclear receptors *Rara, Rxra, Rxrb* or *Rarb,* a classic retinoic acid target gene (Figure 5A). HFD increased serum RBP4 levels in *shScrmbl* control mice compared to chow control and *shHsd17b13* treatment had no effect (Figure 5B). Our previous studies showed that application of the synthetic retinoid Fenretinide can strongly induce retinoic acid responsive nuclear receptor signalling leading to upregulation of classic retinoic acid inducible genes (Morrice *et al.*, 2017, Mcilroy, G. D. *et al.*, 2013). However, our new results suggest that Hsd17b13 does not modulate retinoic acid signalling or retinol homeostasis *in vivo*. To extend these studies further, we examined the effect of Hsd17b13 over-expression and compared it with retinoic acid treatment in cells. Although retinoic acid could increase the expression of *Cyp26a1* and *Rarb*, two classic retinoic acid inducible genes, Hsd17b13 over-expression had no effect alone or in combination with retinoic acid to alter their gene expression (Figure 5C). Hsd17b13 over-expression did increase *Raldh1* expression by >2-fold (p<0.001 by t-test) as did retinoic acid treatment however there was no additive or synergistic interaction (by two-way ANOVA). Our results suggest that Hsd17b13 does not play a significant role in modulating retinoic acid signalling or retinol homeostasis.

### Changing Hsd17b13 levels alters hepatic lipid and phospholipid metabolism

Dis-regulation of a network of hepatic genes involved in fatty acid metabolism in response to over-nutrition contribute to excess hepatic lipid accumulation and thus MASLD. We hypothesised that *Hsd17b13* knockdown mediated decrease in steatotic liver may be associated with a rescue of this impairment. HFD increased hepatic expression of PPARα target genes *Cd36*, *Vldlr, Crat* and *Mogat1* involved in fatty acid transport and metabolism, and *shHsd17b13* treatment inhibited this induction (Figure 6A). To determine if these gene expression changes were due to acute changes in HSD17B13 protein or linked secondarily to decreased hepatic lipid accumulation next, we examined the effect of increased Hsd17b13 expression in cells. We also treated cells with an oleate-rich mixture of fatty-acids containing a low proportion of palmitic acid (oleate/palmitate, 2:1 ratio) representing a cellular model of steatosis that mimics benign chronic steatosis with low toxic and apoptotic effects (Nolan & Larter, 2009, Gomez-Lechon *et al.*, 2007). Increased Hsd17b13 expression increased triglyceride storage in cells (Figure 6B) and increased *Cd36* expression without additional fatty acids in both HepG2 and HEK cells (Figure 6C). Fatty acid treatment further increased triglyceride storage in cells with Hsd17b13 but did not change *Cd36* expression. LXRs promote hepatic de novo lipogenesis and have been linked with Hsd17b13 and MASLD (Su, Wen *et al.*, 2017). LXR agonist treatment did not increase *Cd36* expression and there was no additive or synergistic interaction with Hsd17b13 although LXR stimulation did increase expression of lipogenic target *Fasn* (Supplemental Figure S4).

**Figure 6:**
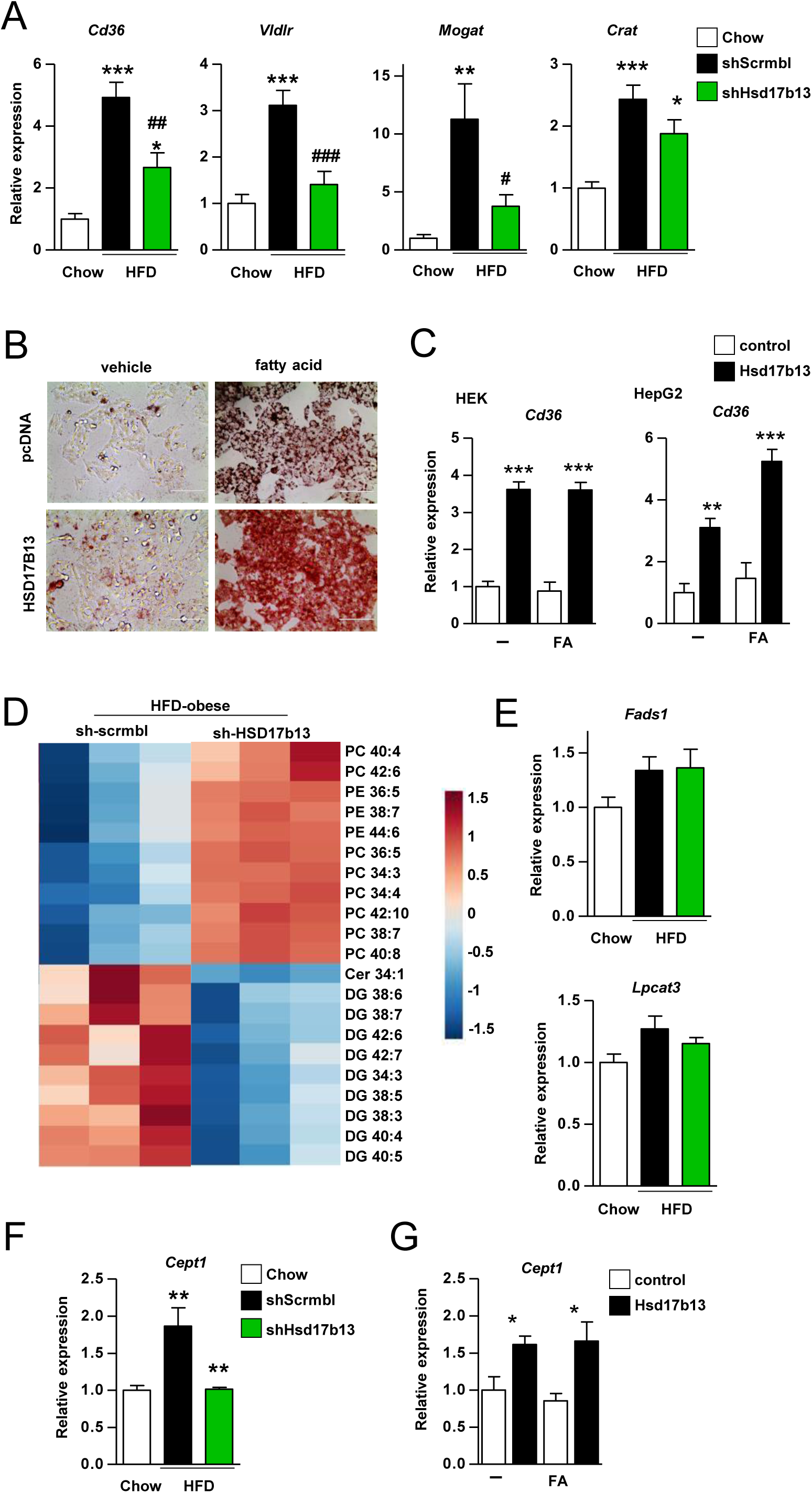
Effect of altered Hsd17b13 levels on lipid and phospholipid metabolism. **A.** Gene expression of hepatic lipid metabolism genes (n=8 per group) in C57BL/6 control CHOW diet mice or HFD obese mice treated with *shHsd17b13* or scrambled shRNA (*shScrmbl*). **B.** HepG2 cells stably expressing HA-tagged Hsd17b13 or pcDNA3.1 control plasmid were treated with 1mM fatty acid (2:1 ratio of oleic and palmitic acid) or methanol (vehicle control). Cells were stained with oil red O. Images are representative of 24 hour treatment. Scale bar = 100µM. **C.** Fatty acid translocase *Cd36* gene expression in HEK293 and HepG2 cells expressing Hsd17b13 or pcDNA3.1 control and treated with unsaturated/saturated fatty acid mixture (2:1 ratio of oleic acid 0.66 mM and palmitic acid 0.33 mM),or methanol (vehicle control). **D.** Lipidomic analysis heatmap of most significant (p<0.05) changes in lipids, enriched in diacylglycerides and phospholipid species in liver tissue from C57BL/6 HFD obese mice treated with *shHsd17b13* or scrambled shRNA (*shScrmbl*), n=3 for each. **E.** and **F**. Gene expression of key regulators of phospholipid metabolism (n=8 per group) **G**. *Cept1* (Choline-enthanolamine phosphotransferase 1) gene expression HEK293 cells expressing Hsd17b13 or pcDNA3.1 control and treated with unsaturated/saturated fatty acid mixture (2:1 ratio of 0.66 mM oleic acid and palmitic acid 0.33 mM) or methanol (vehicle control). All data is presented as mean ± SEM. **A**., **E**. and **F**. were analysed by one-way ANOVA followed by Bonferroni multiple comparison tests where * p≤0.05 , ** p≤0.01 and *** p≤0.001 (HFD groups vs CHOW diet) or # p≤0.05 , ## p≤0.01 and ### p≤0.001 (*shHsd17b13* vs *shscrmbl*). Fads1 and Lpcat3 were increased in HFD, p≤0.05 by t-test. **C**. and **G**. were analysed by two-way ANOVA where * p≤0.05 and *** p≤0.001 (Hsd17b13 cells vs pcDNA control cells), # p≤0.05 and ## p≤0.01 and ### p≤0.001 (fatty acid vs methanol control).

Next, we performed lipidomic analysis of liver tissues to understand the biological function of Hsd17b13 and its role in MASLD. *shHsd17b13* treatment decreased diacylglycerols and increased phospholipids, in particular phosphatidylcholines containing polyunsaturated fatty acids (PUFA) e.g. PC 34:3 and PC 42:10 (Figure 6D). We studied the pathway which is responsible for the production of phospholipids and PUFAs. HFD increased the expression of hepatic *Fads1* and *Lpcat3/Mboat5* (p=0.04 by t-test) and shHsd17b13 treatment did not significantly alter these (Figure 6E, individual data points for mice are in Supplemental Figure S3). HFD markedly increased hepatic *Cept1* expression and *shHsd17b13* treatment normalised *Cept1* expression in mice (Figure 6F). CEPT1 or Choline-enthanolamine phosphotransferase 1 catalyzes the terminal step of the Kennedy pathway and is responsible for phosphatidylcholine (PC) and phosphatidylethanolamine (PE) production (Gibellini & Smith, 2010). It is also reported to produce the endogenous ligand for PPARα, a transcription factor that is a key regulator of hepatic lipid metabolism genes (Chakravarthy *et al.*, 2009). Increased Hsd17b13 expression in cells and increased *Cept1* expression -/+ additional fatty acids (Figure 6G). Thus, Hsd17b13 appears to be a driver of increased hepatic triglyceride storage and phospholipid remodelling, likely via increased *Cept1* and *Cd36* expression, and may explain at least part of the mechanism of it biological function and role in MASLD.

## DISCUSSION

The multiple overlapping secondary pathologies in response to diet-induced obesity such as the development of type 2 diabetes, MASLD, atherosclerosis and CVD are a global medical burden. Currently there is no treatment for MASLD or MASH and thus therapies are urgently needed. We have previously reported the beneficial effects of Fenretinide to prevent the accumulation of hepatic triglycerides via decreased whole-body adiposity and associated insulin resistance. Here our novel RNA-seq study of liver tissue from HFD obese mice -/+ Fenretinide identified major beneficial effects of Fenretinide on hepatic gene expression including key drivers of MASLD e.g. HSD17B13. Thus, we sought to directly knockdown the elevated levels of Hsd17b13 to evaluate its role in driving MASLD and regulation of lipid/phospholipid metabolism.

HSD17B13 gene/protein expression are upregulated in MASLD and transgenic overexpression of Hsd17b13 promotes rapid lipid accumulation in the liver of mice (Su, W. *et al.*, 2014). However, the traditional gene knockout of *Hsd17b13* challenged with several diets to induce steatotic liver and/or fibrosis did not lead to the hypothesised protective phenotype (Ma, Yanling *et al.*, 2021). Our RNAi therapeutic approach, to suppress *Hsd17b13* levels in adult mice with existing HFD-induced liver steatosis to decrease hepatic triglyceride storage is a translational approach to investigate the role of Hsd17b13 in MASLD. This RNAi knockdown strategy has been taken by pharma/biotech in on-going clinical trials e.g. Alnylam Pharmaceuticals and Arrowhead Pharmaceuticals (Amangurbanova *et al.*, 2023). Possible explanations for the discrepancy between the phenotype of mouse genetic knockout and the RNAi knockdown approach, are if Hsd17b13 protein affects other proteins in a mechanism independent of its enzymatic activity or total loss of Hsd17b13 activity is compensated for by another protein/pathway.

Su, Guan and co-workers (Su, Wen *et al.*, 2022) reported that Hsd17b13 interacts with ATGL on lipid droplets to regulate lipolysis in response to cAMP-PKA signalling, a mechanism that resembles a role played by Plin5 (Keenan *et al.*, 2021). Thereby, both Hsd17b13 and Plin5 have been shown to be phosphorylated leading to an increase in ATGL-mediated lipolysis. While this may be beneficial for MASLD, increased hepatic cAMP-PKA signalling is also associated with excess endogenous glucose production via gluconeogenesis in obesity and type 2 diabetes (Yang, H. & Yang, 2016). Thus, it is not clear if this signalling network can be delineated sufficiently to facilitate targeting for the treatment of metabolic diseases. Recently, the X-ray crystal structure of Hsd17b13 has provided insights into a mechanism for association with the lipid droplet and interaction with ATGL or other potential binding partners (Liu *et al.*, 2023). The findings suggest that HSD17B13 may have an important scaffold function at the lipid droplet.

Distinct changes occur in the liver lipidome with MASLD (McGlinchey *et al.*, 2022, Ooi *et al.*, 2021, Velenosi *et al.*, 2022). Moreover, many genes involved in hepatic PUFA and phospholipid metabolism, are important in the formation of biologically important lipids and are linked to metabolic disease by human GWAS and mouse knockout studies e.g. FADS1, ELOVL5, LPCAT3, MBOAT7, CEPT1, *PLA2G4A*/cPLA2α, *PLA2G6*/iPLA2β, PTGDS and PTGES (Zhang, J. Y. *et al.*, 2016, Jalil *et al.*, 2019) .The expression of these genes is also altered by dysregulation of a network of transcription factors such as PPARα, LXR, SREBP. Our lipidomics data suggests that HSD17B13 plays a role in hepatic triglyceride and phospholipid remodelling, whereby knockdown promoted an increase in phosphatidyl-cholines (PC) containing PUFAs eg. PC 34:3 and PC 42:10. Studies of the human genetic variants of Hsd17b13 (protective against severe liver disease) have reported an increase in phospholipids including PC 34:3 (Luukkonen *et al.*, 2020). *shHsd17b13* knockdown normalised liver *Cept1* expression in mice and increased Hsd17b13 expression in cells led to increased *Cept1* expression. These results suggest that Hsd17b13 may regulate phospholipid metabolism and may explain at least part of the mechanism of it biological function and role in MASLD.

Several human genome-wide association studies have identified gene variants in HSD17B13 which generate loss-of-function proteins that associate primarily with protection from severity of MASLD and fibrosis i.e. progression to MASH (Abul-Husn *et al.*, 2018, Anstee *et al.*, 2020, Luukkonen *et al.*, 2020, Ma, Y. *et al.*, 2019, Zhang, H. *et al.*, 2022). In agreement with our *in vivo* data, mouse models have shown that inhibition/knockdown of Hsd17b13 protects against both hepatic triglyceride accumulation and fibrosis (Wang *et al.*, 2022, Su, Wen *et al.*, 2022, Luukkonen *et al.*, 2023). It is well known that feeding mice with special research diets leads to a heterogeneous metabolic and liver disease phenotype that is time dependent and ranges from simple steatosis to more severe cases of steatohepatitis, fibrosis, and expression of fibrosis and inflammatory markers (Loft *et al.*, 2021, Vacca *et al.*, 2024, Clapper *et al.*, 2013).

We did not utilize a specific MASH diet (e.g. GAN diet or choline-deficient HFD) to induce fibrosis or inflammation since these tend to suppress weight gain and adiposity, the primary drivers of MASLD. Moreover, the reciprocal regulation of *Hsd17b13* expression and fibrosis markers was associated with simple HFD-induced obesity +/- beneficial effects of Fenretinide (Figure 1 and 2, Supplemental materials Figure S1).

We determined that *shHsd17b13* knockdown repressed hepatic fatty acid transporter Cd36 expression in mice and reciprocally, increased Hsd17b13 expression in cells led to increased Cd36 expression. Hepatocyte CD36 plays a major role in driving *de novo* lipogenesis and in the development of MASLD (Koonen *et al.*, 2007, Wilson *et al.*, 2016, Rada *et al.*, 2020). Thus, Hsd17b13 may play a role in the regulation of key transcription factors (such as PPARa, LXR and SREBP) which coordinate the expression of hepatic lipid homeostasis genes in response to physiological signals.

These data provide strong evidence for an important role of Hsd17b13 in driving MASLD via regulation of hepatic triglyceride storage and phospholipid metabolism.

Importantly, this beneficial effect is present without an effect on body weight, adiposity or glucose homeostasis suggesting that directly targeting Hsd17b13 is superior to the use of FEN, which has diverse mechanisms of action. In addition, treating MASLD may prevent the development of fibrosis/MASH and create an opportunity to clinically treat these without the need to treat co-morbidities of obesity, type-2 diabetes and cardiovascular diseases. Thus, the translation potential of targeting Hsd17b13 for MASLD/MASH is strong (Zhang, H. *et al.*, 2022, Amangurbanova *et al.*, 2023).

## DATA AVAILABILITY

Data will be made available upon reasonable request. Hepatic RNA-seq data generated in this study (HFD+/-Fenretinide, 20 weeks in C57BL/6J mice) has been deposited in the NCBI Gene Expression Omnibus (GEO) database accession number GSE220684.

## SUPPLEMENTAL MATERIAL

Supplementary Table S1. , Table S2 , Figure S1 (A-E) Figure S2 (A-E) , Figure S3 , Figure S4

## ACKNOWLEDGMENTS

The authors thank the University of Aberdeen (UoA) animal research facility, qPCR core facility (Institute of Medical Sciences, UoA) and Linda Davidson (NHS Grampian) for their technical contributions regarding animal studies, qPCR and histology respectively. RNA-sequencing was carried out at the Centre for Genome Enabled Biology and Medicine (CGEBM, UoA); thanks to Diane Stewart, Sophie Shaw and Elaina Collie-Duguid for support with sample prep, bioinformatics reporting and project supervision. Thanks to Lars Grøntved (University of Southern Denmark) for invaluable discussions about the RNA-seq bioinformatic analysis. Lipidomics was performed at Glasgow Polyomics (University of Glasgow). The authors also wish to thank Mirela Delibegovic (UoA) for invaluable discussions throughout the study and review of the manuscript and members of the Delibegovic-Mody joint lab (especially Joe Smith, Sarah Kamli-Salino and Abby Mitchell) for their technical contributions to experiments and analysis of data.

## DISCLAIMERS

None

## AUTHOR CONTRIBUTIONS

GDM and N.Mody conceived and designed research; S.Mahmood, N. Morrice, DT, S. Milanizadeh, SW and PDW performed experiments and analyzed data; JJR and N.Mody interpreted results of experiments; S.Mahmood, S. Milanizadeh, SW, DT and PDW prepared figures; S.Mahmood drafted the manuscript and N.Mody edited and revised the manuscript, S.Mahmood, N. Morrice, GDM and N.Mody approved final version of manuscript.

