## Supplemental for "Hydroxysteroid 17-beta dehydrogenase 13 *(Hsd17b13)* knockdown attenuates liver steatosis in high-fat diet obese mice"

Wed 26th Jun 2024 main text and suppl. data revised for submission to J. Physiol.

### Supplementary Materials

**Supplementary Table S1**
**Supplementary Table S2**

**Supplementary Figure S1 (A-E)**
**Supplementary Figure S2 (A-E)**
**Supplementary Figure S3**
**Supplementary Figure S4**

**Supplementary table S1: Summary of metabolic parameters in C57BL/6 mice fed HFD +/- FEN for 20 weeks or *db/db* +/- FEN for 14 weeks.**

| | | | |
|---|---|---|---|
| **HFD +/- FEN diet study (20 weeks)** | **Chow** | **HFD** | **FEN-HFD** |
| Final body weight (g) | 31.8 ± 0.6 | 45.2 ± 0.87 * | 39.8 ± 1.5 *, # |
| Adiposity at 12 weeks (fat mass in g) | 4.6 ± 0.2 | 13.4 ± 0.5 * | 10.3 ± 0.9 *, # |
| Serum glucose at 12 weeks (mmol/L) | 11.9 ± 0.7 | 20.9 ± 0.2 * | 11.6 ± 0.6 # |
| Serum insulin (ng/mL) | 0.7 ± 0.01 | 2.8 ± 0.3 * | 1.6 ± 0.2 * |
| Liver triglyceride (µg/mg) | 15.4 ± 2.0 | 56.5 ± 5.0 *** | 29.4 ± 5.7 ## |
| Serum FGF21 (ng/mL) | 0.4 ± 0.05 | 3.5 ± 1.2 * | 0.3 ± 0.07 # |
| ***db/db* +/- FEN study (14 weeks)** | **Lean** | ***db/db*** | **FEN-*db/db*** |
| Final body weight (g) | 32.8 ± 0.6 | 44.8 ± 1.5 *** | 50.9 ± 2.0 ***, ## |
| Serum glucose at 4 weeks (mmol/L) | 8.8 ± 0.2 | 29.9 ± 1.4 *** | 16.8 ± 2.9 *, ### |
| Serum insulin (ng/mL) | 1.0 ± 0.1 | 3.6 ± 1.2 * | 11.5 ± 2.6 *, # |
| Liver triglyceride (µg/mg) | 8.3 ± 0.9 | 53.8 ± 5.4 * | 62.0 ± 9.7 * |
| Serum FGF21 (ng/mL) | 0.7 ± 0.16 | 2.7 ± 0.2 * | 0.8 ± 0.23 # |

Data was previously reported in :

**Mcilroy GD**, Delibegovic M, Owen C, Stoney PN, Shearer KD, McCaffery PJ, Mody N: Fenretinide treatment prevents diet-induced obesity in association with major alterations in retinoid homeostatic gene expression in adipose, liver, and hypothalamus. *Diabetes.* 62:825-836, **2013**

**Morrice N**, Mcilroy GD, Tammireddy SR, Reekie J, Shearer KD, Doherty MK, Delibegovic M, Whitfield PD, Mody N: Elevated fibroblast growth factor 21 (FGF21) in obese, insulin resistant states is normalised by the synthetic retinoid fenretinide in mice. *Sci Rep.* 7:43782, **2017**.

Data are shown as mean ± SEM and significance was determined by one-way ANOVA followed by *post-hoc* tests. Differences are marked * p<0.05, ** p<0.01 *** p<0.001 vs chow or lean (HFD or *db/db* control mice respectively); # p<0.05 ## p<0.01 ### p<0.001 vs HFD or *db/db*.

**Supplementary table S2: List of primers**

| **Mouse qPCR (liver tissue)** | | |
|---|---|---|
| **Gene** | Forward | Reverse |
| *Avpr1a* | GCTGGCGGTGATTTTCGTG | GCAAACACCTGCAAGTGCT |
| *Cd68* | TGTCTGATCTTGCTAGGACCG | GAGAGTAACGGCCTTTTTGTGA |
| *Cd36* | AAGTAGCACAGGAGCCTCAG | TGCTGTTCTTTGCCACGTCA |
| *Cept1* | AGTCTTCTACTGCCCTACAGC | TCCAAGAACCACAAAAACTGTCG |
| *Col1a* | CCAAAGGTGCTGATGGTTCT | ACCAGCTTCACCCTTGTCAC |
| *Crat* | CATCCGTCAGCCTCCATAG | CTCGGATGGCCCGGTCAG |
| *Cyp46a1* | AGCCGCTATGAGCACATCC | CCATACTTCTTAGCCCAATCCAG |
| *Fads1* | CCAGCTTTGAACCCACCAA | CATGAGGCCCATTCGCTCTA |
| *Gstm2* | ACACCCGCATACAGTTGGC | TGCTTGCCCAGAAACTCAGAG |
| *Hsd17b2* | ACCTTGTTCCTCTTATCCGTGG | ACCGAAACCGGAATCAGCAC |
| *Hsd17b13* | ATTCCCCGGAGAAGGAAATCT | CAGCCTGCCTATTCCGTGT |
| *Lpcat3* | GGCCTCTCAATTGCTTATTTCA | AGCACGACACATAGCAAGGA |
| *Mogat* | TGGACGCCAGTTTGGTTCCAG | TGCTCTGAGGTCGGGTTCA |
| *Nono (housekeeping reference gene)* | GCCAGAATGAAGGCTTGACTAT | TATCAGGGGGAAGATTGCCCA |
| *Rara* | CGC CAA GGG AGC TGA ACG GG | GGG TGG CTG GGC TGC TTC TG |
| *Rarß* | CGCGAGCCCTTCCTCCTGC | AAAAGCCCTTGCACCCCTCGC |
| *Rdh16* | TTGTGACAGCCACTCAGTGGGT | CCAGTACATGTGCAAAGTCCTGC |
| *Rxra* | ATGGACACCAAACATTTCCTGC | CCAGTGGAGAGCCGATTCC |
| *Rxrß* | CCACCTCTTACCCCTTCAGC | TGGAAGAACTGATGACTGGGA |
| *Tgfß* | ATCCTUTCAAACTAAGGCTCG | ACCTCTTTAGCATAGGTAGTCCGC |
| *Timp1* | GCAACTCGGACCCTGGTCATAA | CGCCCCGTGATGAGAACT |
| *Timp2* | TCAGAGCCAAAGCAGTGAGC | GCCGTGTAGATAAACTCGATGTC |
| *Vldlr* | GGCAGCAGGCAATGCAATG | GGGCTCGTCACTCCAGTCT |

| **Human qPCR (HEK, HepG2 cells)** | | |
|---|---|---|
| *CYP26A1* | CTCACATTGACAGGGATTGGA | GCTGGCCAGTGGACCGACAC |
| *HPRT (housekeeping reference gene)* | CATTATGCTGAGGATTTGGAAAG | CTACAATGTGATGGCCTCCCA |
| *HSD17B13* | ATCACAAAAGCACTTCTTCCATC | AGACCTCTGTGAAAGCCAAC |
| *RALDH1* | TGTTAGCTGATGCCGACTTG | TTCTTAGCCCGCTCAACACT |
| *RARb* | TCTCAGTGCCATCTGCTTAATCTG | CCAGCAATGGTTCTTGTAGCTTATC |

**Supplementary Figure S1**

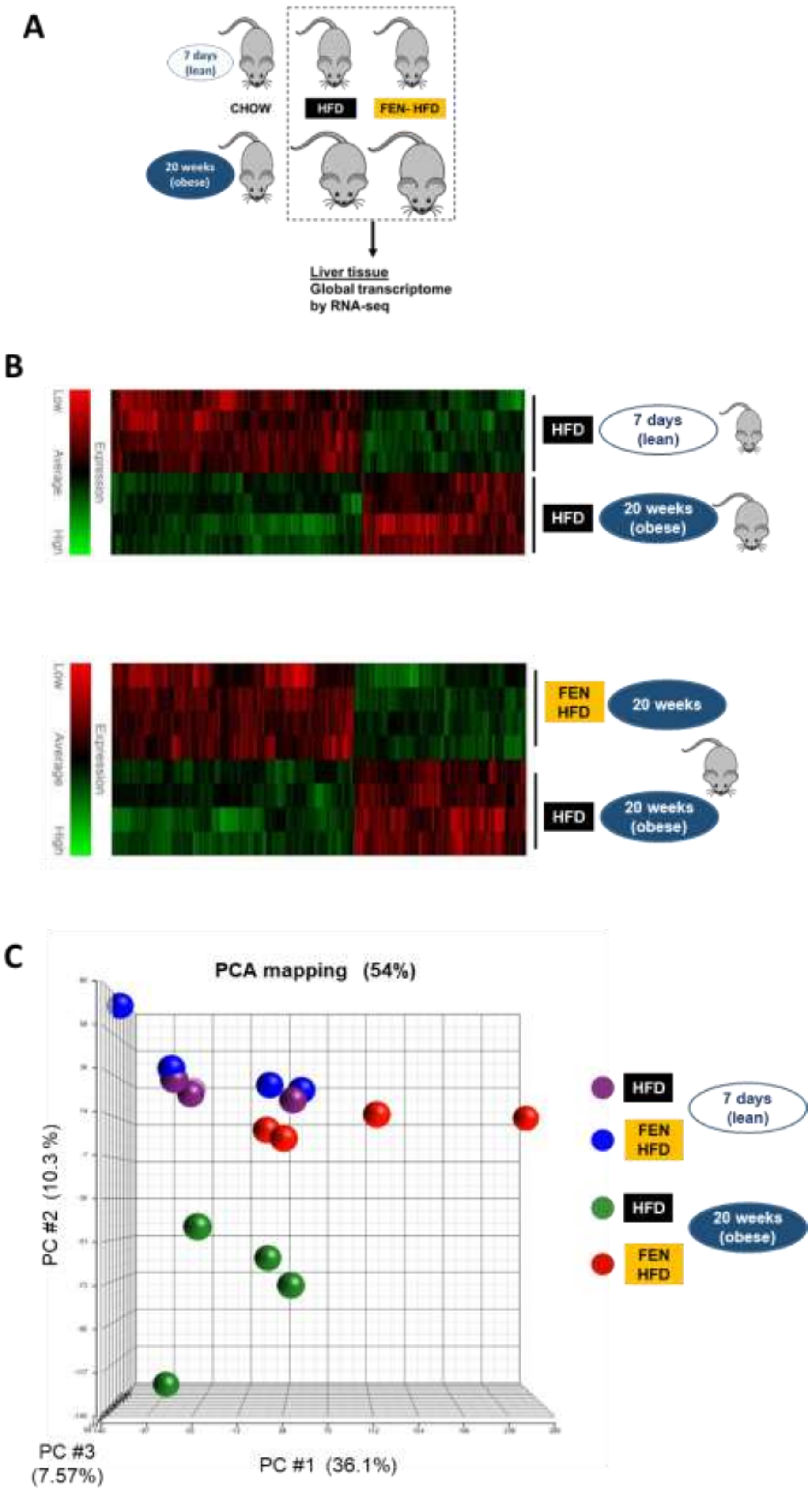

**Supplementary Figure S1**

**D**

| Term | Fold enrichment | Normalised p value | Number of genes |
|---|---|---|---|
| Triglyceride biosynthetic process | 150.37 | $1.01e^{-4}$ | 4 |
| Fatty acid catabolic process | 146.61 | $7.04e^{-3}$ | 3 |
| CDP-diacylglycerol biosynthetic process | 112.76 | $9.11e^{-3}$ | 3 |
| Very long-chain fatty acid metabolic process | 91.63 | $1.23e^{-3}$ | 3 |
| Fatty acid beta-oxidation | 55.53 | $1.05e^{-4}$ | 5 |
| Fatty acid metabolic process | 43.86 | $2.09e^{-16}$ | 14 |
| Phospholipid biosynthetic process | 33.13 | $7.67e^{-3}$ | 4 |
| Lipid metabolic process | 24.49 | $3.21e^{-24}$ | 23 |
| Metabolic process | 13.72 | $1.22e^{-8}$ | 13 |

### Supplementary Figure S1

# E

| | FEN-HFD 20 wks | | HFD 20 wks | | HFD 20 wks | | FEN-HFD 20 wks | | FEN-HFD 20 wks | | |
|---|---|---|---|---|---|---|---|---|---|---|---|
| | vs HFD 7 days | | vs HFD 7 days | | vs FEN HFD 7 days | | vs FEN HFD 7 days | | vs HFD 20 wks | | |
| | Fold Change | padj | Fold Change | padj | Fold Change | padj | Fold Change | padj | Fold Change | padj | |
| *Mogat1* | 4.1 | *** | 14.0 | # | 15.2 | # | 4.3 | *** | -2.7 | ** | *Mogat1* |
| *Agpat9* | 4.1 | # | 6.1 | # | 3.8 | # | 2.5 | *** | -1.4 | * | *Agpat9* |
| *Crat* | 2.1 | ** | 5.1 | # | 5.8 | # | 2.4 | # | -2.4 | # | *Crat* |
| *Vldlr* | 1.3 | ns | 3.6 | *** | 5.2 | *** | 1.8 | * | -2.4 | ** | *Vldlr* |
| *Cyp46a1* | 1.6 | ns | 5.1 | # | 4.0 | *** | 1.3 | ns | -3.0 | ** | *Cyp46a1* |
| *Rdh16* | 1.2 | ns | 3.2 | # | 3.3 | # | 1.3 | ns | -2.5 | *** | *Rdh16* |
| *Gstm2* | 1.6 | * | 3.6 | # | 3.9 | # | 1.7 | * | -2.1 | ** | *Gstm2* |
| *Pex11a* | 2.0 | ** | 3.3 | # | 2.6 | ** | 1.5 | ns | -1.6 | *** | *Pex11a* |
| *Vnn1* | 2.8 | # | 4.6 | # | 3.2 | # | 1.9 | ** | -1.6 | *** | *Vnn1* |
| *Abcc3* | 1.7 | *** | 3.0 | # | 2.2 | *** | 1.2 | ns | -1.8 | ** | *Abcc3* |
| *Pparg* | 2.6 | ** | 3.3 | *** | NaN | NaN | NaN | NaN | -1.2 | ns | *Pparg* |
| *Insig2* | 1.2 | NaN | 2.1 | ** | 2.0 | ** | NaN | NaN | NaN | NaN | *Insig2* |
| *Cd36* | 2.1 | *** | NaN | NaN | NaN | NaN | 2.5 | # | NaN | NaN | *Cd36* |
| *Acaa1b* | 1.4 | ns | 2.6 | # | 1.7 | *** | NaN | NaN | -1.8 | * | *Acaa1b* |
| Acot4 | 2.6 | ** | 4.3 | # | NaN | NaN | 1.9 | * | -1.6 | * | Acot4 |
| Abcb1a | 2.0 | ** | 3.2 | *** | NaN | NaN | 2.3 | ** | -1.6 | * | Abcb1a |
| *Gpam* | 0.9 | ns | 1.8 | * | 2.4 | ** | 1.2 | ns | -2.0 | ** | *Gpam* |
| *Plin5* | 1.1 | ns | 1.7 | ** | NaN | NaN | 1.1 | ns | -1.5 | * | *Plin5* |
| *Gm15441* | NaN | NaN | 2.9 | ** | 2.2 | * | NaN | NaN | -2.1 | * | *Gm15441* |
| *Pklr* | 1.0 | ns | 2.0 | ** | 2.4 | # | 1.2 | ns | -2.0 | ** | *Pklr* |
| *Osbpl5* | 0.9 | ns | 2.1 | ** | 2.9 | *** | 1.2 | ns | -2.3 | ** | *Osbpl5* |
| *Ralgps2* | 0.7 | * | 1.3 | ns | 1.7 | ** | 0.9 | ns | -1.9 | *** | *Ralgps2* |
| *Pnldc1* | 2.8 | ** | 7.7 | # | 6.9 | # | 2.6 | ** | -2.1 | * | *Pnldc1* |
| *Nek2* | 1.0 | ns | 2.0 | * | 2.1 | ** | 1.0 | ns | -2.1 | ** | *Nek2* |
| *Adamtsl2* | NaN | NaN | NaN | NaN | 2.0 | ** | 0.7 | ns | -2.7 | ** | *Adamtsl2* |
| *Lpar1* | 0.9 | ns | 2.0 | ** | 2.3 | ** | 1.0 | ns | -2.2 | ** | *Lpar1* |
| *Tm6sf2* | 1.0 | ns | 1.6 | ** | 2.3 | *** | 1.5 | ns | -1.5 | * | *Tm6sf2* |
| *Pnpla3* | 1.6 | ns | 2.4 | * | 2.8 | NaN | 1.8 | * | -1.7 | ns | *Pnpla3* |
| *Hsd17b13* | 0.7 | * | 1.5 | * | 2.1 | NaN | 0.9 | ns | -2.3 | ** | *Hsd17b13* |
| *Inhbe* | 1.2 | ns | 1.6 | * | 1.9 | ** | 1.5 | ns | -1.3 | ns | *Inhbe* |
| *Slc39a5* | 2.4 | * | 5.5 | *** | 4.2 | *** | 1.8 | ns | -2.0 | * | *Slc39a5* |
| *Slc16a5* | 2.3 | ** | NaN | NaN | 4.6 | *** | 1.9 | * | -2.4 | ** | *Slc16a5* |
| *Slc17a4* | 0.8 | ns | 1.8 | * | NaN | NaN | NaN | NaN | -2.2 | ** | *Slc17a4* |
| *Slc16a13* | 2.5 | ** | 2.7 | ** | 2.5 | ** | 2.3 | ** | -1.0 | ns | *Slc16a13* |
| *Timp1* | NaN | NaN | 3.2 | * | NaN | NaN | 30.9 | ns | -1.9 | ns | *Timp1* |
| *Timp2* | 0.8 | ns | 1.4 | * | 1.7 | ** | 1.0 | ns | -1.7 | ** | *Timp2* |
| *Col1a1* | NaN | NaN | 2.2 | ** | NaN | NaN | NaN | NaN | -1.8 | ns | *Col1a1* |
| *Wwtr1* | 1.7 | ** | 1.9 | # | 1.6 | *** | 1.5 | ** | -1.1 | ns | *Wwtr1* |
| *Ctgf* | 1.3 | ns | 2.4 | ** | NaN | NaN | 1.2 | ns | -1.8 | ** | *Ctgf* |
| *Col15a1* | 0.9 | ns | 1.7 | * | 2.8 | *** | 1.4 | ns | -2.0 | ** | *Col15a1* |
| *Col12a1* | 1.3 | ns | 2.4 | ** | 2.3 | ** | 1.2 | ns | -1.8 | ** | *Col12a1* |
| *Col20a1* | 1.2 | ns | 0.6 | * | 0.6 | * | 1.2 | ns | -0.5 | ** | *Col20a1* |
| *Col8a1* | 0.5 | ns | 1.4 | ns | 1.9 | * | 0.6 | ns | -3.1 | ** | *Col8a1* |
| *Col16a1* | 0.8 | ns | 1.5 | ns | 2.0 | ** | 1.0 | ns | -1.9 | ** | *Col16a1* |
| *Fads1* | 0.8 | NaN | NaN | NaN | 2.8 | # | 1.2 | ns | -2.2 | *** | *Fads1* |
| *Napepld* | 1.5 | ns | 2.5 | ** | 2.5 | * | 1.3 | ns | -1.5 | ns | *Napepld* |
| *Ptgfr* | NaN | NaN | 2.2 | * | 2.2 | * | NaN | NaN | -2.1 | * | *Ptgfr* |
| *Ptges* | 3.3 | ** | 2.1 | * | 2.1 | ns | 2.4 | ** | -0.6 | ns | *Ptges* |
| *Pctp* | 1.7 | ** | 2.0 | ** | 2.0 | ns | 1.1 | ns | -1.2 | ns | *Pctp* |
| *Pla2g6* | 1.5 | * | 1.8 | ** | 1.8 | ns | 1.2 | ns | -1.2 | ns | *Pla2g6* |
| *Ptgds* | -4.7 | *** | 0.4 | * | 0.4 | ns | NaN | NaN | -1.8 | ns | *Ptgds* |

**Supplementary Figure S1: Differential gene expression from RNA-seq data.**

(A) Schematic diagram of RNA-seq from liver tissue from C57BL/6J mice fed HFD+/-FEN for 20 weeks (obese) or HFD+/-FEN for 7 days (lean).

(B) Heatmap of differentially expressed genes in HFD 20 weeks (obese) vs HFD 7 days (lean), *upper panel*. Heatmap of differentially expressed genes in FEN-HFD 20 weeks vs HFD 20 weeks (obese), *lower panel*.

(C) Principle Component Analysis (PCA) of differentially expressed genes from all four groups HFD+/-FEN for 20 weeks (obese) or HFD+/-FEN for 7 days (lean) demonstrating HFD 20 weeks (obese) mice having a gene expression profile most different to the other groups. Moreover, the FEN-HFD 20 weeks mice having a gene expression profile more similar to both lean groups (HFD+/-FEN for 7 days), than the HFD 20 weeks (obese) group. Plot was generated using Partek Genomics Suite.

(D) Gene ontology biological component categories of genes differentially expressed with HFD 20 weeks (obese) mice and normalised by Fenretinide treatment FEN-HFD 20 weeks mice. Performed using David v6.8.

(E) Selection of reciprocally regulated liver genes in HFD (obese)+/-FEN, involved in triglyceride synthesis and fatty acid/LC-PUFA metabolism and genes associated with MASLD/MASH. Differentially expressed gene analysis from all four groups HFD +/- FEN for 20 weeks (obese) vs HFD +/- FEN for 7 days (lean) and rescue in FEN-HFD 20 weeks vs HFD 20 weeks (obese). RNA sequencing of total RNA extracted from frozen liver, as previously described (10) . Full data generated in this study has been deposited in the NCBI Gene Expression Omnibus (GEO) database accession number GSE220684. Values have been presented as Fold Change and significance, adjusted p-value # $<10^{-15}$ , *** $<10^{-8}$ , ** $<10^{-3}$ , * <0.05, ns not significant. NaN, no data available.

Wed 26th Jun 2024 main text and suppl. data revised for submission to J. Physiol.

### Supplementary Figure S2

A

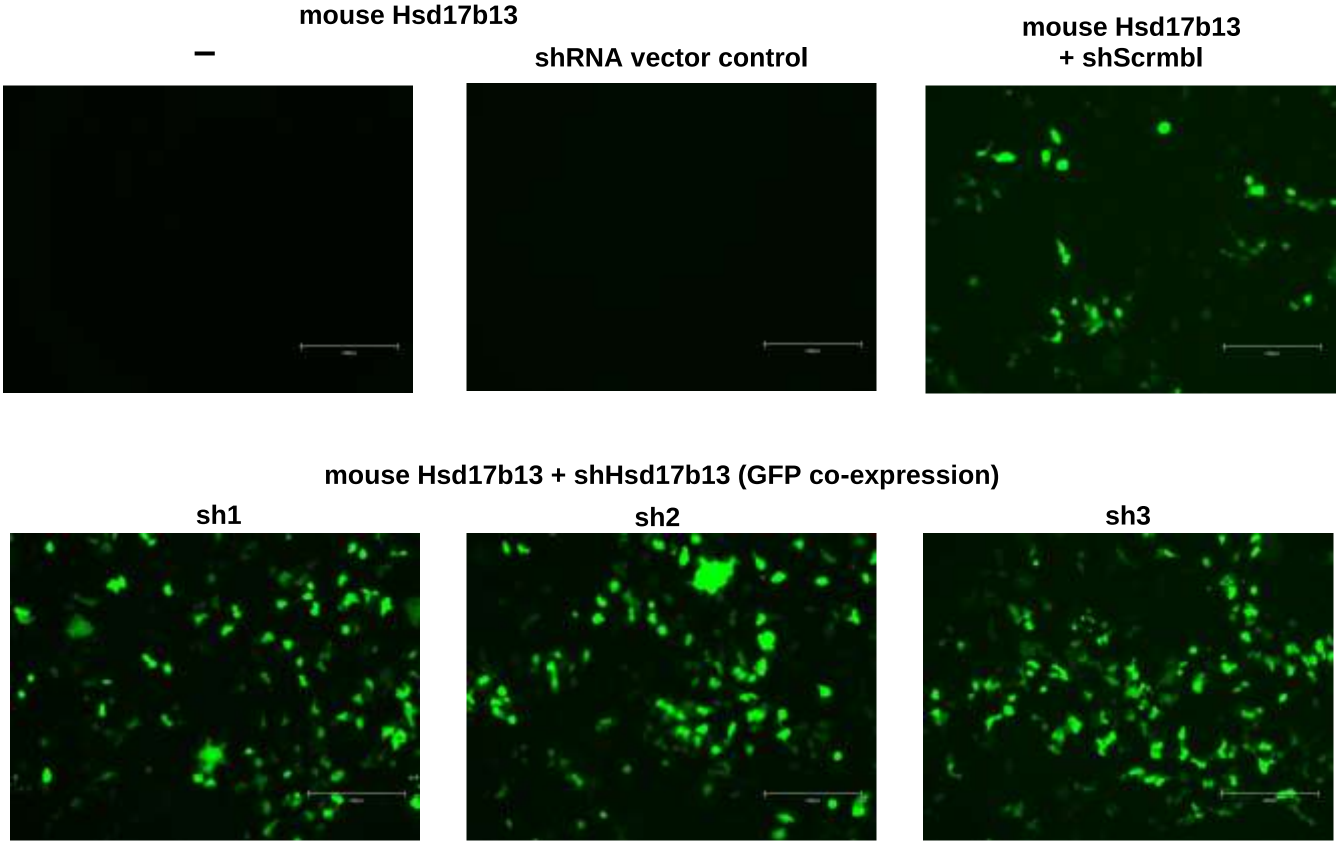

B

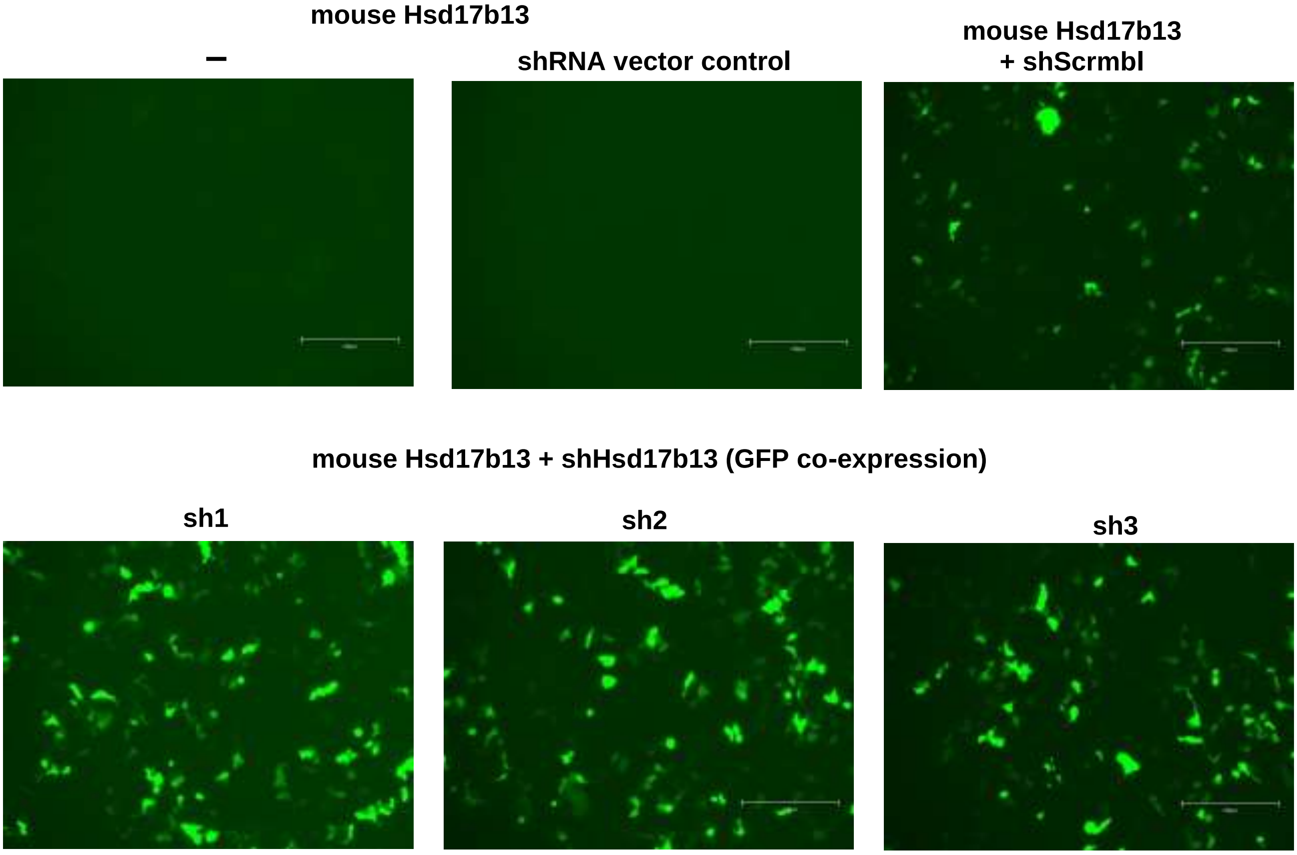

**Supplementary Figure S2**

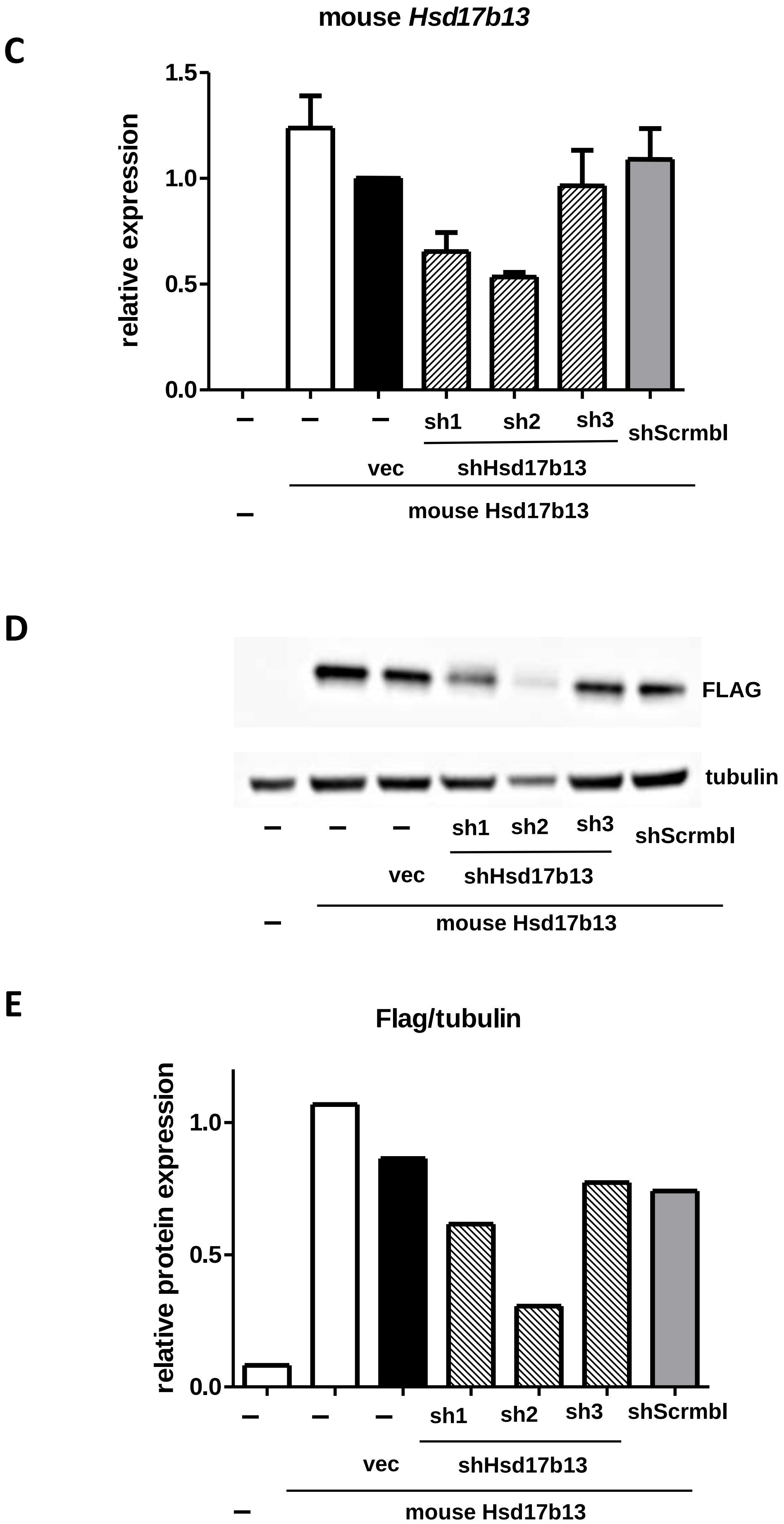

**Supplementary Figure S2: Testing shRNA for knockdown efficiency of mouse *Hsd17b13* overexpressed in Human Embryonic Kidney (HEK)-293 cells.**

Mouse *Hsd17b13* was overexpressed in HEK293 cells (*white bar*) by transient transfection (0.25µg plasmid DNA, 24 hours), followed by transfection with shRNA vector control (*black bar*), one of three candidate shRNA (sh1-3) (*hatched bars*) or scrambled control (shScrmbl) (*grey bar*) (0.75 µg plasmid DNA, 48 hours).

(A-E) Representative data shown from n=3 biological replicates with similar results.
(A-B) GFP (co-expression with shRNA expression) fluorescence microscopy images,
(C) Mouse *Hsd17b13* gene expression (determined by qPCR),
(D-E) Mouse *Hsd17b13* protein expression (determined by FLAG-tag immunoblot). Tubulin was used as a housekeeping/loading control. Blots were quantified using ImageJ software (E).

Scale bars are 300 µm in fluorescent images. Data shown is gene or protein expression fold change compared to mouse *Hsd17b13* with shRNA vector control (black bar). Human *HPRT* was used as housekeeping control gene to normalise gene expression using the Pfaffl method.

**Supplementary Figure S3**

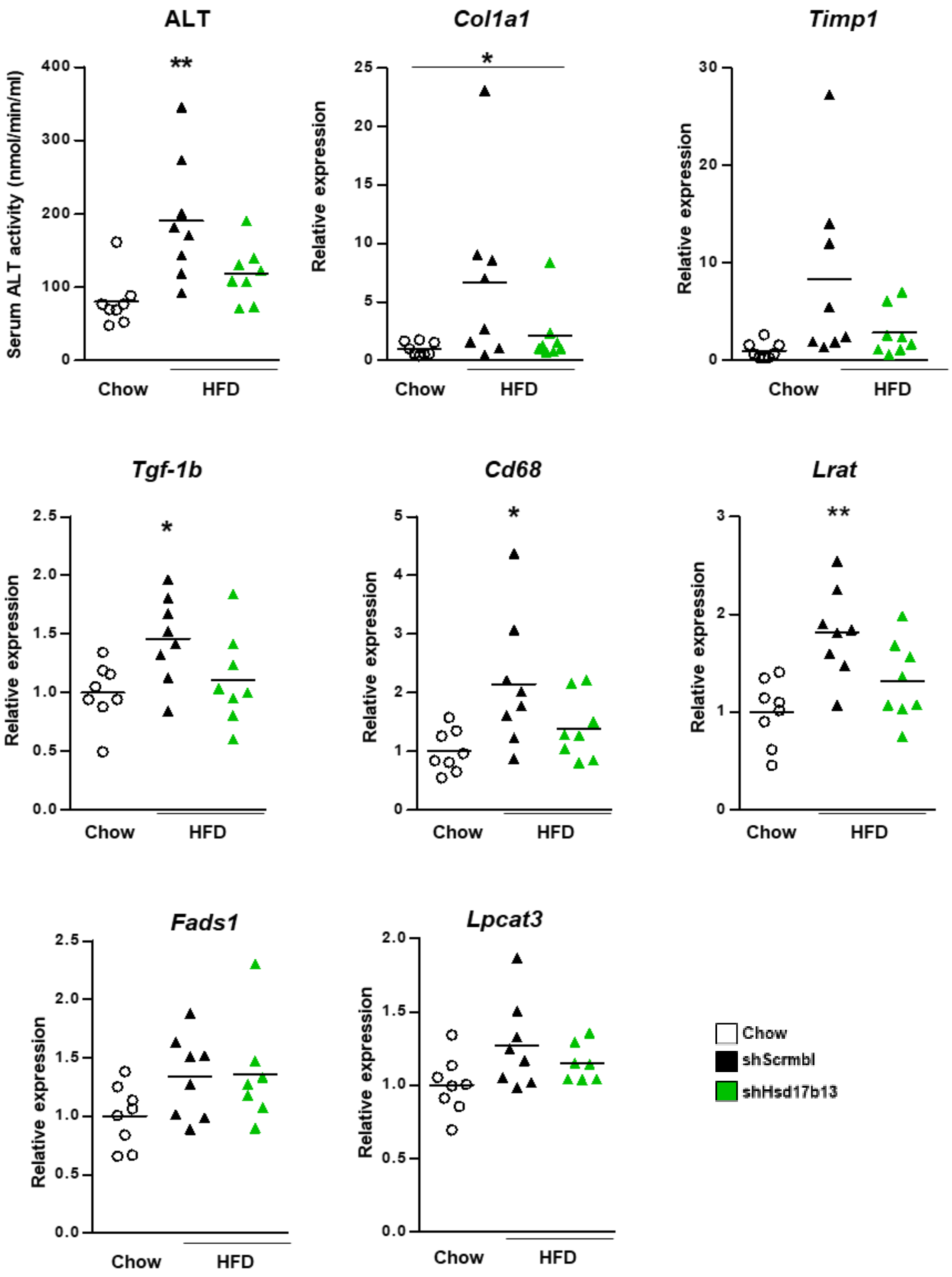

**Supplementary Figure S3 - Individual data points for *mice C57BL/6 control CHOW diet or HFD obese treated with shHsd17b13 or scrambled control (shScrmbl)* in graphs presented in Figure 4, Figure 5 and Figure 6**. Serum ALT activity and gene expression of fibrosis markers in liver tissue, *Collagen type I alpha 1* (*Col1a1*), *Collagen type IV alpha 1* (*Col4a1*), *Tissue inhibitor of metalloproteinases* (*Timp*)*-1* and *Timp2, Transforming growth factor beta* (*Tgf-β*)*-1* and Cluster of Differentiation (*Cd*)*-68, lecithin-retinol acyltransferase* (*Lrat*), key regulators of phospholipid metabolism *Fatty acid desaturase 1* (*Fads1*) and *lysophosphatidylcholine acyltransferases* (*Lpcat*)-*3.* Data are presented as individual data points and mean, and analysed by one-way ANOVA followed by Bonferroni multiple comparison tests where * $p \le 0.05$, ** $p \le 0.01$ and *** $p \le 0.001$ (comparing HFD groups to control CHOW diet) or # $p \le 0.05$ and ## $p \le 0.01$ (comparing HFD-shHsd17b13 to HFD-shscrmbl).

**Supplementary Figure S4**

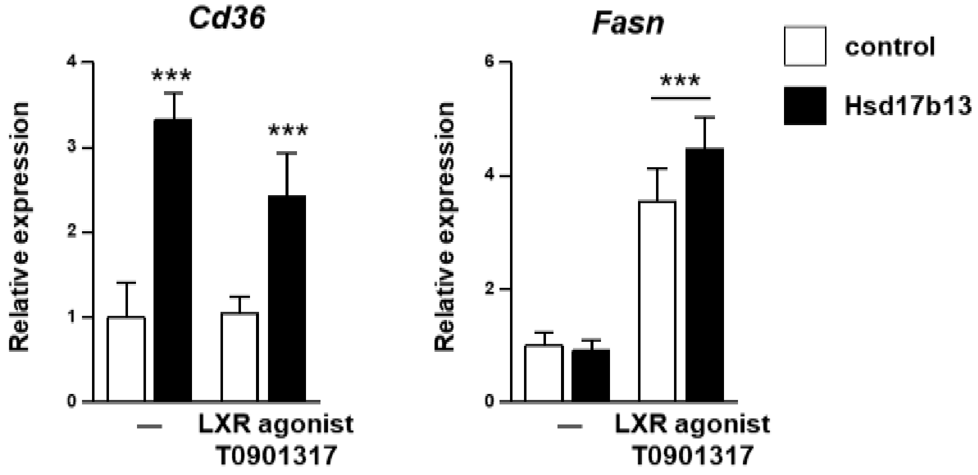

**Supplementary Figure S4 – Effect of altered Hsd17b13 levels with LXR agonist T0901317 stimulation on gene expression of lipid metabolism regulators.**

Relative mRNA expression of *Cd36* and *Fasn* in HepG2 cells stably expressing HA-tagged Hsd17b13 or pcDNA3.1 control plasmid. Cells were treated with 1μM LXR agonist T0901317 or DMSO (vehicle control) for 24 hours. For each group n = 2 biological repeats (4 technical). Significance was calculated by two-way ANOVA with Tukey's post hoc test, P value <0.05. There was no significant interaction between Hsd17b13 expression and T0901317 stimulation.
